# Exploring the druggability of the UEV domain of human TSG101 in search for broad-spectrum antivirals

**DOI:** 10.1101/2024.11.15.623737

**Authors:** Fernando Montero, Marisa Parra-López, Alejandro Rodríguez-Martínez, Javier Murciano-Calles, Pedro Buzon, Ziying Han, L.-Y. Lin, Maria C. Ramos, Javier Ruiz-Sanz, Jose C. Martinez, Marco Radi, Christiane Moog, Sandra Diederich, Ronald N. Harty, Horacio Pérez-Sánchez, Francisca Vicente, Francisco Castillo, Irene Luque

## Abstract

The ubiquitin E2 variant domain of TSG101 (TSG101-UEV) plays a pivotal role in protein sorting and virus budding by recognizing PTAP motifs within ubiquitinated proteins. Disruption of TSG101-UEV/PTAP interactions has emerged as a promising strategy for the development of host-oriented broad-spectrum antivirals with low susceptibility to resistance. TSG101 is a challenging target characterized by an extended and flat binding interface, low affinity for PTAP ligands, and complex binding energetics. Here, we assess the druggability of the TSG101-UEV/PTAP binding interface by searching for drug-like inhibitors and evaluating their ability to block PTAP recognition, impair budding, and inhibit viral proliferation. A discovery workflow was established combining in vitro miniaturized HTS assays and a set of cell-based activity assays including high-content bimolecular complementation, virus-like particle release measurement, and antiviral testing in live virus infection. This approach has allowed us to identify a set of chemically diverse molecules that block TSG101-UEV/PTAP binding with IC50s in the low μM range, and able to disrupt the interaction between full-length TSG101 and viral proteins in human cells and inhibit viral replication. State-of-the-art molecular docking studies reveal that the active compounds exploit binding hotspots at the PTAP binding site, unlocking the full binding potential of the TSG101-UEV binding pockets. These inhibitors represent promising hits for the development of novel broad-spectrum antivirals through targeted optimization and are also valuable tools for investigating the involvement of ESCRT in the proliferation of different virus families and study the secondary effects induced by the disruption of ESCRT/virus interactions.

**Importance:** Many viruses rely on the interaction between TSG101 and viral proteins containing PTAP motifs for their proliferation. Here we show that these interactions can be efficiently blocked by drug-like compounds that impair budding and replication of viruses from different families. We have also provided valuable insights into the determinants of high affinity for these small molecule inhibitors that open new avenues for developing the identified candidates into broad-spectrum antivirals with low susceptibility to resistance.

## 1. Introduction

Human Tumor-susceptibility Gene 101 (TSG101) is a member of the class E family of vacuolar protein-sorting proteins and an essential component of the Endosomal Sorting Complex Required for Transport (ESCRT) pathway, a conserved complex of about 30 proteins in charge of driving membrane remodeling and fission in multiple cellular processes, including cytokinesis, autophagy, multivesicular body generation, extracellular vesicle biogenesis, as well as plasma, nuclear and endolysosomal membrane repair. ESCRT proteins are organized in four main complexes (ESCRT-0, -I, -II, and -III), which are sequentially recruited to the target membrane and assisted by accessory proteins NEDD4 (Neuronal precursor cell-expressed developmentally down-regulated protein 4) and ALIX (ALG-2 interacting protein X) that provide selectivity in protein recruitment (Vietri et al., 2020; Calistri et al., 2021; Lemus and Goder, 2022). TSG101 controls ESCRT-I formation and orchestrates early events of membrane trafficking and scission through the interaction with a PTAP motif in Hepatocyte Growth Factor-Regulated Tyrosine Kinase Substrate (HRS) within the ESCRT-0 complex (Lu et al., 2003) and the recognition of ubiquitinated cargo, playing, thus, a key scaffolding role for the subsequent engagement of ESCRT-II and ESCRT-III.

Due to its functional versatility, the ESCRT machinery is sequestered by many intracellular pathogens for replication, assembly, or egress (Calistri et al., 2021; Meng and Lever, 2021; Rivera-Cuevas and Carruthers, 2023). ESCRT recruitment is typically mediated by small peptide sequences called viral Late domains (L-domains) found in the Gag polyproteins of retroviruses and in the structural proteins of many families of enveloped RNA viruses (Ahmed et al., 2019; Ferraiuolo et al., 2020; Rose, 2021). Three types of L-domains have been identified containing PT/SAP, YPX_n_L/LxxLF, or PPxY conserved motifs that act as cellular adaptors engaging ESCRT functions through specific interactions with TSG101, ALIX, and NEDD4, respectively (Freed, 2002; Bieniasz, 2006; Welker et al., 2021). These interactions are essential to complete the replication cycle of viruses that bud from the plasma membrane (filoviruses, arenaviruses, rhabdoviruses alphaviruses, and paramyxoviruses) or from intracellular membranes, such as the ER, the Golgi apparatus, or the ER-Golgi compartment (coronaviruses, flaviviruses, and bunyaviruses, among others) (Welker et al., 2021; Rivera-Cuevas and Carruthers, 2023). Molecules disrupting ESCRT/L-domain interactions have been shown to block the egress of different viruses (Tavassoli et al., 2008; Han et al., 2014; Loughran et al., 2016; Lennard et al., 2019; Han et al., 2021; Castillo et al., 2022) and constitute potential host-oriented antivirals with a wide spectrum of action and low susceptibility to the development of resistance. Thus, as one of the main factors responsible for pathogenic ESCRT recruitment, TSG101 is an attractive target for therapeutic intervention.

PTAP L-domains engage TSG101 through its Ubiquitin-conjugating enzyme E2 Variant (UEV) domain, mimicking TSG101/HRS interactions. TSG101-UEV shares the E2 ligase typical fold, although it contains an additional N-terminal α-helix, presents an extended β-hairpin (tongue), and lacks the two C-terminal helices, exposing a hydrophobic groove that contains the PTAP binding site (Pornillos et al., 2002a; Pornillos et al., 2002b; Palencia et al., 2006). TSG101-UEV binds PTAP-containing cellular targets with low binding affinity (Kd ∼ 200-300 μM for HRS). It also binds ubiquitin (Kd ∼ 500-800 μM) but is catalytically inactive due to the absence of the active-site cysteine, which is replaced by a tyrosine residue (Y_110_) in the TSG101 sequence (Garrus et al., 2001; Katzmann et al., 2001; Sundquist et al., 2004). The ubiquitin binding site lies at a concave region at the lower half of the β-sheet, flanked by the β4-α3 loop at the vestigial active site (lip), and the β-hairpin tongue (Pornillos et al., 2002a; Sundquist et al., 2004). The different location of the two interaction sites allows the simultaneous or even cooperative binding of ubiquitin and PTAP-containing proteins (Sundquist et al., 2004; Strickland et al., 2022).

Blocking TSG101-UEV/PTAP interactions is a promising strategy for the development of novel antivirals with a very wide spectrum of action and low susceptibility to resistance. However, TSG101 is a challenging target. As other proline-recognition modules, it presents extended and relatively flat interfaces, low affinity, and complex binding energetics elicited by the interplay of conformational contributions, water-mediated interactions, and strong enthalpy/entropy compensation effects (Ferreon and Hilser, 2003; Palencia et al., 2010; Martin-Garcia et al., 2012; Iglesias-Bexiga et al., 2019). Nonetheless, TSG101-UEV/PTAP binding is somewhat “hot-spot” in nature, being dominated by the PTAP core, which establishes a distinctive and highly conserved pattern of direct and water-mediated interactions (Im et al., 2010; Murciano-Calles et al., 2024). Most of the binding energy is contributed by the second proline in the motif (P_0_) (Aasland et al., 2002) that packs tightly into a hydrophobic pocket (xP pocket) shared by the different families of proline-recognition modules, including SH3, WW, EVH, or GYF domains (Ball et al., 2005). Binding affinity and specificity are modulated by residues flanking the PTAP_0_ core, with positions +2/+3 being key for tight binding. We have recently shown that optimized packing and electrostatic interactions implicating residues at these positions elicit one-to-two order of magnitude improvements in affinity, being responsible for the tight binding of the HIV-1 and Ebola virus (EBOV) L-domain sequences (Kd∼20-50 μM) and the low micromolar affinity of optimized phage display peptides (Kd∼1-2 μM) (Murciano-Calles et al., 2024). These highly localized hot-spots make the TSG101-UEV/PTAP interface amenable to inhibition by drug-like compounds, as illustrated by previous attempts that have produced peptidomimetics (Liu et al., 2008; Kim et al., 2011), cyclic peptides (Tavassoli et al., 2008; Lennard et al., 2019), and small-molecule compounds (Liu et al., 2011; Lu et al., 2014) able to block PTAP recognition and, in some cases, inhibit budding in simplified virus systems.

In this work, we set out to further assess the druggability of the TSG101-UEV/PTAP binding interface by searching for drug-like inhibitors with different chemical scaffolds and evaluating their ability to block PTAP recognition, impair budding, and inhibit viral proliferation. For this, we have set up a discovery workflow including in vitro Differential Scanning Fluorescence and AlphaScreen miniaturized HTS assays combined with cell-based assays to assess inhibition that include high-content bimolecular complementation, measurement of virus-like particle release, and antiviral testing in live virus infection. This approach has allowed us to identify a set of chemically diverse molecules blocking TSG101-UEV/L-domain binding with IC50s in the low μM range, some of which efficiently disrupt the interaction between full-length TSG101 and L-domain-containing viral proteins in human cells and show antiviral activity. Our study confirms that targeting the PTAP binding site in TSG101-UEV is an efficient and innovative approach to the search for host-oriented antivirals. The identified compounds could constitute valuable starting points to develop novel antivirals with a broad spectrum of action and effective tools for early intervention against emerging pathogens or bioterrorism attacks with modified viruses. These molecules will also be of interest to assess the extent and mechanisms of toxicity and gauge the secondary effects associated with the disruption of ESCRT/virus interactions, and to further our understanding of the implication of ESCRT in the proliferation of different virus families.

## 2. Results

### 2.1. High Throughput Screening of small-molecule libraries against TSG101-UEV

A Differential Scanning Fluorimetry (DSF) assay was set up and optimized in a 96-well format for the screening of small-molecule libraries against the UEV domain of TSG101. Thermal denaturation profiles of TSG101-UEV at different protein concentrations confirmed that under assay conditions TSG101-UEV unfolds as a monomeric protein producing a single and highly reproducible cooperative transition (Figure S1A). TSG101-UEV also shows good tolerance to DMSO, remaining correctly folded and thermally stable at DMSO concentrations up to 10% (Figure S1B). The PD1 (GRVVE**PSAP**PWEGP) and PD2 (YGGSLV**PSAP**PMPPM) peptides, previously identified in our laboratory by phage display and binding to TSG101-UEV with dissociation constants in the low micromolar range (Murciano-Calles et al., 2024), were used as positive controls. These peptides induce significant Tm shifts compared to the free TSG101-UEV protein (negative control), confirming that the DSF assay is robust and presents adequate performance and resolution (Ź values between 0.5-0.7) (Table 1).

**Table 1.** Quality parameters for the DSF screening assay.

| Control | Kd (25°C)<br>(μM) | Tm<sup>1</sup><br>(°C) | Tm shift<br>(°C) | Average<br>Z' factor | S/N |
| --- | --- | --- | --- | --- | --- |
| TSG101-UEV | - | 63.0±0.4 | - | - |  |
| PD-1 | 2.2 | 66.7±0.3 | 3.7 | 0.5 | 9.8 |
| PD-2 | 1.7 | 68.8±0.2 | 5.8 | 0.7 | 15.3 |

<sup>1</sup>Average values ± standard deviation

A total of 8753 compounds from the NCI Diversity Set IV + Natural Products Set III libraries (National Institutes of Health, USA) and the ASINEX Protein-Protein Interaction library were screened in a 96-well format for binding to 2 µM TSG101-UEV in PBS, pH 7.5, 8% DMSO at a final concentration of 800 µM (Figure 1). These collections include small molecules containing pharmacologically desirable features as well as larger natural products and macrocyclic derivatives designed as secondary structure or epitope mimetics. They cover a wide range of structural and chemical diversity with good physicochemical properties (see Figure S2 for details). A total of 56 compounds (16 NCI & 40 ASINEX) stabilizing the protein over 1°C in the primary DSF screen were selected for further analysis.

**Figure 1.**
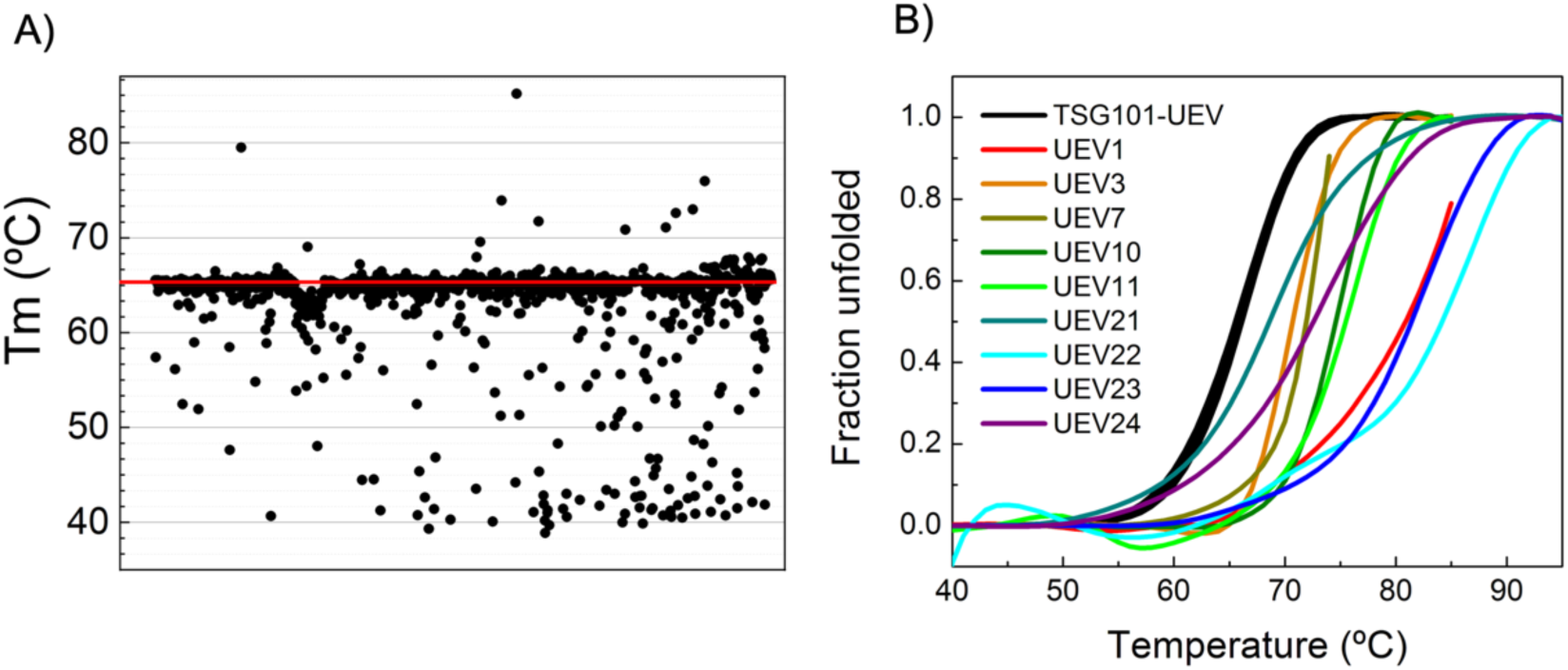
Primary high-throughput screening using differential scanning fluorimetry to identify binders of TSG101-UEV. **(A)** General results of the screening of the NCI Diversity Set IV and the Natural Products Set III libraries against TSG101-UEV. **(B)** Normalized DSF profiles for TSG101-UEV in the presence of 9 hits selected for further characterization (ΔTm > 1 °C). Profiles were obtained in PBS, 8% DMSO at 2 µΜ TSG101-UEV and 800 µM compound. Seven replicas of the denaturation profile for the free TSG101-UEV domain were used as negative controls (black lines).

The ability of the selected compounds to bind TSG101-UEV and to disrupt its interactions with the HIV1 p6-Gag protein (HIV1-p6), containing a PTAP L-domain plus a YPX_n_L motif targeting ALIX-V, was validated using an orthogonal secondary assay based on the AlphaScreen (Amplified Luminiscence Proximity Homogeneous Assay, Revvity) technology. This is a competition binding assay in which the disruption of a pre-formed TSG101-UEV/HIV1-p6 complex (Kd = 27 µM) (Garrus et al., 2001) produces a luminescence change. Cherry-picking assays were performed in triplicate at a fixed 400 µM compound concentration. For the confirmed positives, dose-response experiments were carried out and IC50 values determined (Figure 2 and Table 2).

**Figure 2.**
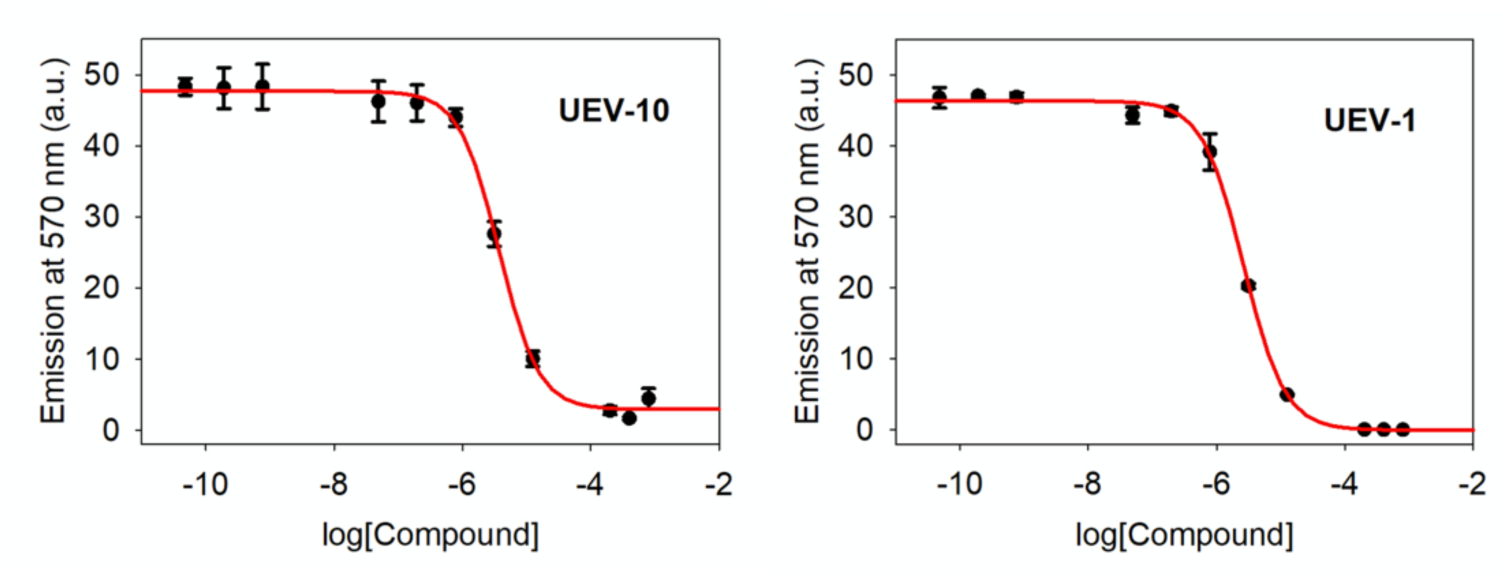
Dose-response curves obtained using AlphaScreen competition assays against the TSG101-UEV/HIV1-p6 pre-formed complex. Symbols represent the average of three independent experiments and error bars show the standard deviation at a 95% confidence interval. The red line corresponds to the best fit of results to the IC50 fitting equation (see Materials and Methods).

**Table 2.** Characterization of inhibitors of TSG101-UEV/HIV1-p6 interactions.

| Compound | $\Delta T_m$<br>(°C) | AlphaScreen<br>signal reduction<sup>1</sup> | IC<sub>50</sub><sup>2</sup><br>(μM) | K<sub>d</sub><br>(μM) | qAC<sub>50</sub><sup>3</sup><br>(μM) |
| --- | --- | --- | --- | --- | --- |
| UEV-1 | 14.6 | 100% | 2.6 ± 0.4 | 2.4 | > 50 |
| UEV-3 | 4.7 | 61% | 150 ± 40 | 140 | > 50 |
| UEV-7 | 6.0 | 58% | 280 ± 40 | 260 | > 50 |
| UEV-10 | 8.1 | 97% | 3.7 ± 0.6 | 3.4 | 21.6 ± 0.4 |
| UEV-11 | 11.1 | 100% | 5.3 ± 0.3 | 4.9 | 7.4 ± 0.9 |
| UEV-21 | 4.0 | 34% | 1 ± 2 | 1 | > 50 |
| UEV-22 | 22.8 | 40% | 5 ± 2 | 5 | 34.3 ± 0.7 |
| UEV-23 | 16.8 | 41% | 5 ± 1 | 4 | nd<sup>4</sup> |
| UEV-24 | 7.9 | 33% | 4 ± 1 | 4 | 43 ± 2 |

<sup>1</sup>Cherry picking measurements at 400 μM compound concentration. <sup>2</sup>IC<sub>50</sub> values derived by AlphaScreen competitive assay. <sup>3</sup>Half-maximal activity values derived from the hill equation analysis of MTT assays in HEPG2 cells. Errors correspond to the SE values derived from triplicate experiments. <sup>4</sup>Not determined.

A total of nine compounds were validated as ligands of TSG101-UEV able to disrupt TSG101-UEV/HIV1-p6 binding with IC50 values in the low-mid micromolar range. The structures and predicted properties of the selected compounds are detailed in Table S1. Inspection of their chemical structures reveal two distinct groups of inhibitors: a) a set of small and rigid molecules (250-400 Da and < 4 rotatable bonds) rich in N-substituted cycles that induce a complete loss of the AlphaScreen signal at saturating concentrations and steep IC50 curves (compounds UEV1 to UEV-11) and b) a set of larger and more flexible tetracyclic heterocycles containing a tetrahydropyran rings fused with a piperidine linked to aromatic amino acids (compounds UEV21 to UEV24). Even though the latter compounds have low micromolar IC50 values, they elicit more gradual curves and a partial disruption of the AlphaScreen signal.

Overall, these TSG101-UEV inhibitors present good predicted physicochemical and drug-like properties (Table S2), complying with the Lipinsky and Veber rules and show TPSA values that correlate well with passive molecular transport through membranes (Ertl et al., 2000; Lipinski et al., 2001). Most of them are predicted to be moderately soluble with good lipophilicity. Nonetheless, some (UEV-1 and UEV-10) include Pan Assay Interference Structures (PAINS) (Baell and Holloway, 2010) and others (UEV-1, -3, -10, -11, and -21) bear unwanted groups according to Brenk’s alerts (Brenk et al., 2008) that could be targeted for removal in future med-chem optimization. Also, these molecules were predicted to have high gastro-intestinal absorption, although there is a discrepancy between algorithms with respect to their ability to pass the blood-brain barrier (BBB). No significant inhibition of cytochrome P4502D6 and cytochrome P4503A4 was predicted (Table S3).

The selected TSG101-UEV/HIV1-p6 inhibitors showed low toxicity on standard MTT assays in human HEPG2 cell lines, except for compound UEV-11 that was found to be toxic at concentrations above 10 μM (Table 2 and Figure S3). These results confirm that, despite the challenging features of the TSG101-UEV/PTAP interaction, low-micromolar drug-like molecules interfering with the recognition PTAP L-domain peptides by the TSG101-UEV domain can be identified.

### 2.2. Bimolecular complementation assay (BiMC) between full-length TSG101 and viral L-domain containing proteins

The ability of the selected compounds to enter human cells and inhibit the interaction between the TSG101 and HIV1-Gag proteins in a biologically relevant context was assessed using a well-established YFP-based bimolecular complementation assay (Liu et al., 2011) coupled to a high-content image analysis system. Co-expression of the NYFP-TSG101 (full length TSG101 fused to the N-terminal domain of YFP) and CYFP-HIV1-Gag (full-length Gag polyprotein from HIV-1 fused to the C-terminal domain of YFP) constructs efficiently reconstitutes YFP producing a strong fluorescent signal that is lost upon disruption of the TSG101/HIV1-Gag interaction by an effective inhibitor. The inhibitory activity of the nine selected compounds was quantified as the reduction in the total YFP signal compared with control wells treated with 0.5% DMSO. To assess to what extent the reduction in fluorescence signal is due to the specific disruption of the TSG101/HIV1-Gag interaction or to unspecific toxicity, a high-content analysis of the total cell counts based on Hoechst nuclear staining was used to quantify cell viability. The ability of the inhibitors to disrupt TSG101/PTAP interactions within different contexts was explored using an equivalent BiMC system targeting the interaction between TSG101 and the VP40 matrix protein from EBOV (NYFP-TSG101/CYFP-EBOV-VP40). EBOV-VP40 contains at its N-terminus a PPxY L-domain overlapping the PTAP core (_5_ILPTAPPEY_13_) as well as an ALIX-binding YPX_n_L L-domain.

The different compounds were assayed in triplicate against the HIV1 and EBOV BiMC systems at different inhibitor concentrations. Three experimentally independent replicas were performed, producing a total of 9 measurements per compound and concentration to provide statistically robust values. As illustrated in Figure 3, all compounds produced very similar results for the two BiMC systems. Compound UEV-10 showed good dose-dependent inhibitory activity, efficiently blocking TSG101 interactions at 10 μM in the context of the two BIMC systems (HIV1-Gag and EBOV-VP40) without significant toxicity (Figure S4) and maintaining significant inhibitory levels at lower concentrations. At 10 μM UEV-11 elicits an 80% reduction in the YFP signal with high toxicity, as expected from the MTT results. Nonetheless, the βYFP-signal/toxicity ratio of 5 suggest some level of specific inhibition (Figure S5) that is confirmed at lower concentrations (UEV-11 retains a 35-50% inhibition with less than 10% loss of cellular viability at 0.1 μM) (Figure S5). Other N-substituted compounds (UEV-1 and UEV-7) as well as the UEV-21 and UEV-24 tetracyclic inhibitors produced less significant results, eliciting a 30% reduction in the YFP signal at 10 μM concentration. The effects of compounds UEV-22 and UEV-23, showing solubility issues and irreproducible results (data not shown), could not be properly measured in these assays.

**Figure 3.**
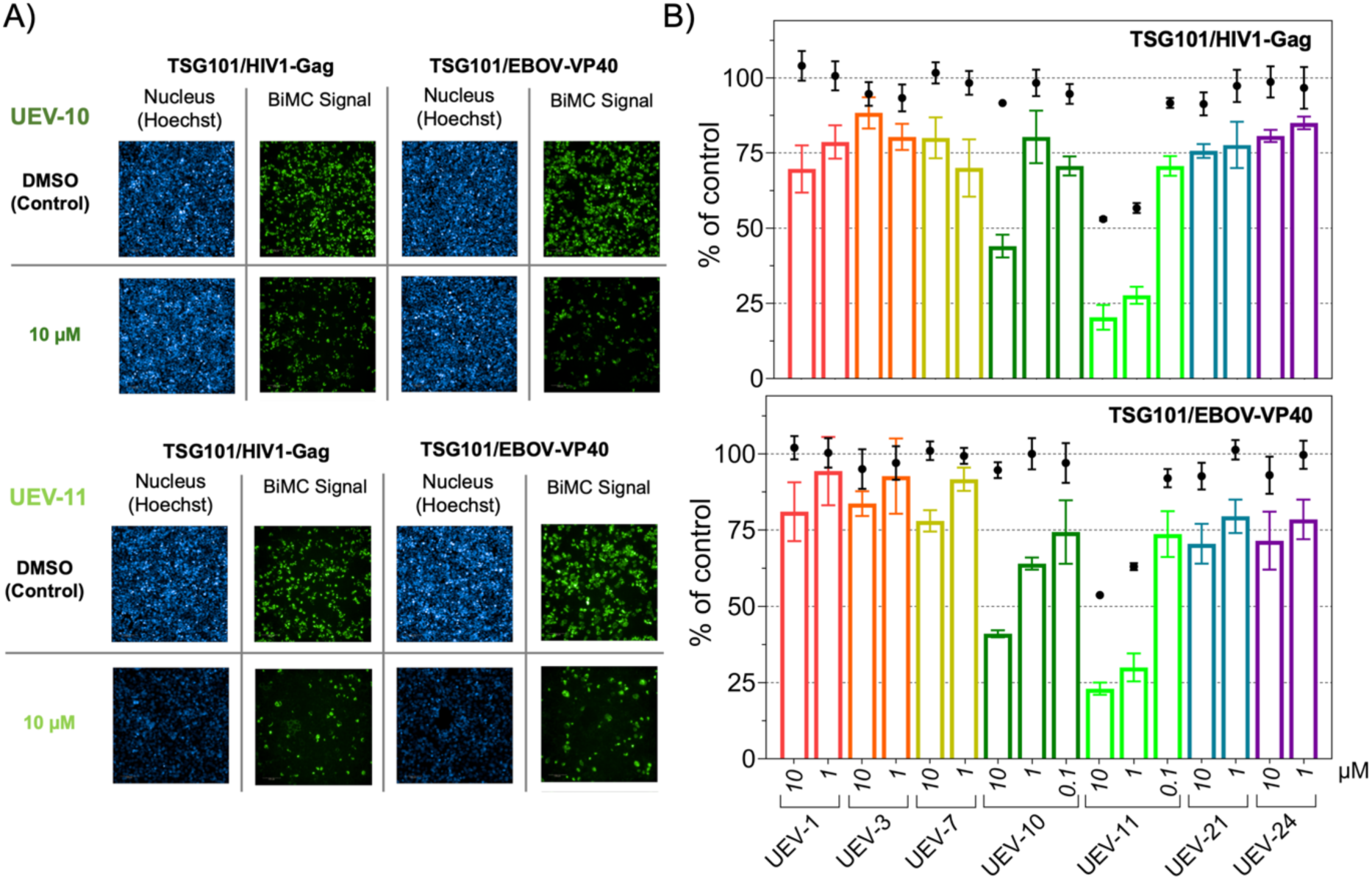
Disruption of the protein-protein interaction between full-length TSG101 and PTAP-containing viral proteins followed by a YFP-based BiMC assay. **(A)** Representative images of HEK293T cells co-expressing NYFP-TSG101 and CYFP-EBOV-VP40 or CYFP-HIV1-Gag fusion proteins in the absence (control) and presence of different concentrations of UEV-10 (dark green) and UEV-11 (light green) compounds. The blue signal corresponds to the Hoechst-stained cell nuclei reporting on the number of viable cells in the well and green signal correspond to the YFP fluorescence. **(B)** Effect of different inhibitors on the YFP signal corresponding to the NYFP-TSG101/CYFP-HIV1-Gag (upper panel) or NYFP-TSG101/CYFP-EBOV-VP40 (lower panel) interactions. The percentage of the YFP signal retained in the presence of varying concentrations of the selected compounds compared to the control (no inhibitor, 0.5% DMSO) is shown as colored bars. The corresponding cell viability values, calculated as the percentage of total cells compared to controls are shown as black dots. Error bars show the standard error of the mean. All reported values correspond to the average of a total of 9 measurements obtained from three independent replicas performed in triplicate. The YFP signal and the number of viable cells were quantified using the Harmony software (Revvity).

### 2.3. Antiviral activity in cell-based assays

The capacity of the TSG101-UEV inhibitors to impair HIV-1 replication was evaluated using a variety of cell-based assays in human cells. No inhibitory effect was detected after one round of infection with pseudoviruses (data not shown), as could be expected considering that these compounds have been selected to target the budding process which is one of the very last stages of viral replication. To assess the effect of the different compounds on the infectivity of the newly produced viruses, multiple rounds HIV-1 infection were analyzed by measurement of Renilla luciferase (RLU) produced by a replication-competent infectious molecular clone (IMC) encoding a subtype C env gene (BJOX) and the Renilla luciferase gene (Sarzotti-Kelsoe et al., 2014). Alternatively, HIV SF162 replication was detected by measurement of intracellular p24-HIV core protein by Flow cytometry (Klingler et al., 2022). Human lymphoblastoid cell lines CEM-SS and A3CCR5 or primary peripheral blood mononuclear cells 5PBMC were used. The toxicity of the inhibitors was assessed in parallel to virus inhibition using the Renilla-Glo® Luciferase Assay System and the Selective Index (SI: ratio HIV-1 inhibition/toxicity) was calculated.

As summarized in Table 3, several compounds inhibited HIV-1 replication, although some toxicity was observed at high compound concentrations resulting in decreased cell counts. A positive SI was, nonetheless, observed for some virus/cell combinations (shown in bold in Table 3). UEV 22 and 24 showed a positive SI in both assays, and the inhibition was confirmed on primary PBMC corresponding to the N-substituted compounds with binding affinities in the low micromolar range (Table 2) as well as the UEV-22 and UEV-24 inhibitors. These results show that UEV-22 and UEV-24 demonstrating high binding affinity to TSG101-UEV were able to decrease the infectivity of new viruses released after multiple rounds of infection at concentrations that do not affect cell proliferation.

**Table 3.** HIV-1 inhibition assays.

| Compound | RLU |  | SF162 + flow cytometry |  |  |
| --- | --- | --- | --- | --- | --- |
|  | CEM cells | A3R5,7 cells | CEM cells | A3R5,7 cells | PBMC |
| | IC50/Tox (μM)<br>SI | IC50/Tox (μM)<br>SI | IC50/Tox (μM)<br>SI | IC50/Tox (μM)<br>SI | IC50/Tox (μM)<br>SI |
| UEV-1 | 281/151<br>**0.5** | 96.4/186<br>**1.9** | 53/96<br>**1.8** | 97/280<br>**2.9** | 75/100<br>**1.3** |
| UEV-3 |  | 93.6/>100<br>nd<sup>1</sup> | >100/>100<br>nd<sup>1</sup> | >100/>100<br>nd<sup>1</sup> | 100/>100<br>nd<sup>1</sup> |
| UEV-7 |  | 94.4/>100<br>nd<sup>1</sup> | 75/>100<br>nd<sup>1</sup> | >100/>100<br>nd<sup>1</sup> | 100/>100<br>nd<sup>1</sup> |
| UEV-10 | 31/18<br>**0.6** | 12.5/24<br>**1.9** | 67/89<br>**1.3** | 98/98<br>**1** | 17/27<br>**1.6** |
| UEV-11 | 0.2/1.9<br>**9.7** | 0.6/1.3<br>**2.3** | 1/<1<br>nd | 6/<1<br>nd | 6/<1<br>nd |
| UEV-21 |  | 73.1/>100<br>nd<sup>1</sup> | 81/>100<br>nd<sup>1</sup> | 100/>100<br>nd<sup>1</sup> | 100/>100<br>nd<sup>1</sup> |
| UEV-22 |  | 15.7/71<br>**4.5** | 11/22<br>**2.1** | 21/29<br>**1.4** | 61/91<br>**1.5** |
| UEV-24 |  | 46.1/>100<br>**>2** | 13/37<br>**2.7** | 61/87<br>**1.4** | 76/102<br>**1.3** |

<sup>1</sup> SI values were not calculated for compound/cell combinations for which toxicity was not observed under the conditions studied (Tox > 100 μM).

To test whether the TSG101-UEV inhibitors also had an inhibitory effect against live EBOV, Vero E6 cells were infected with a recombinant EBOV expressing GFP (rgEBOV-VP30-GFP) in the presence of TSG101-UEV inhibitors. At 72 h post infection, GFP expression was monitored as a read out for EBOV replication. As shown in Figure 4A, addition of UEV-10 in increasing concentrations up to 10 µM led to a clear reduction in GFP expression and thus, EBOV replication. However, the addition of UEV-10 at 10 µM had some cell toxic effect. Analysis of virus titers in the supernatant demonstrated a 2-log reduction of virus titer in the presence of 7 µM UEV-10 without toxicity (Figure 4B). A slightly stronger reduction of virus titer was observed for incubation with 10 µM UEV-10, however, at this concentration cell toxic effects might have an additional negative impact. Due to high toxicity, UEV-11 could only be assayed at concentrations below 3 µM. Under these conditions, incubation in the presence of UEV-11 only had a minor effect on EBOV spread and no significant reduction in virus titer was detected for these inhibitors. No significant inhibition of EBOV replication was observed for the rest of the inhibitors tested (Figure S6), including compounds UEV-22 and UEV-24 that had demonstrated antiviral activity against HIV-1.

**Figure 4.**
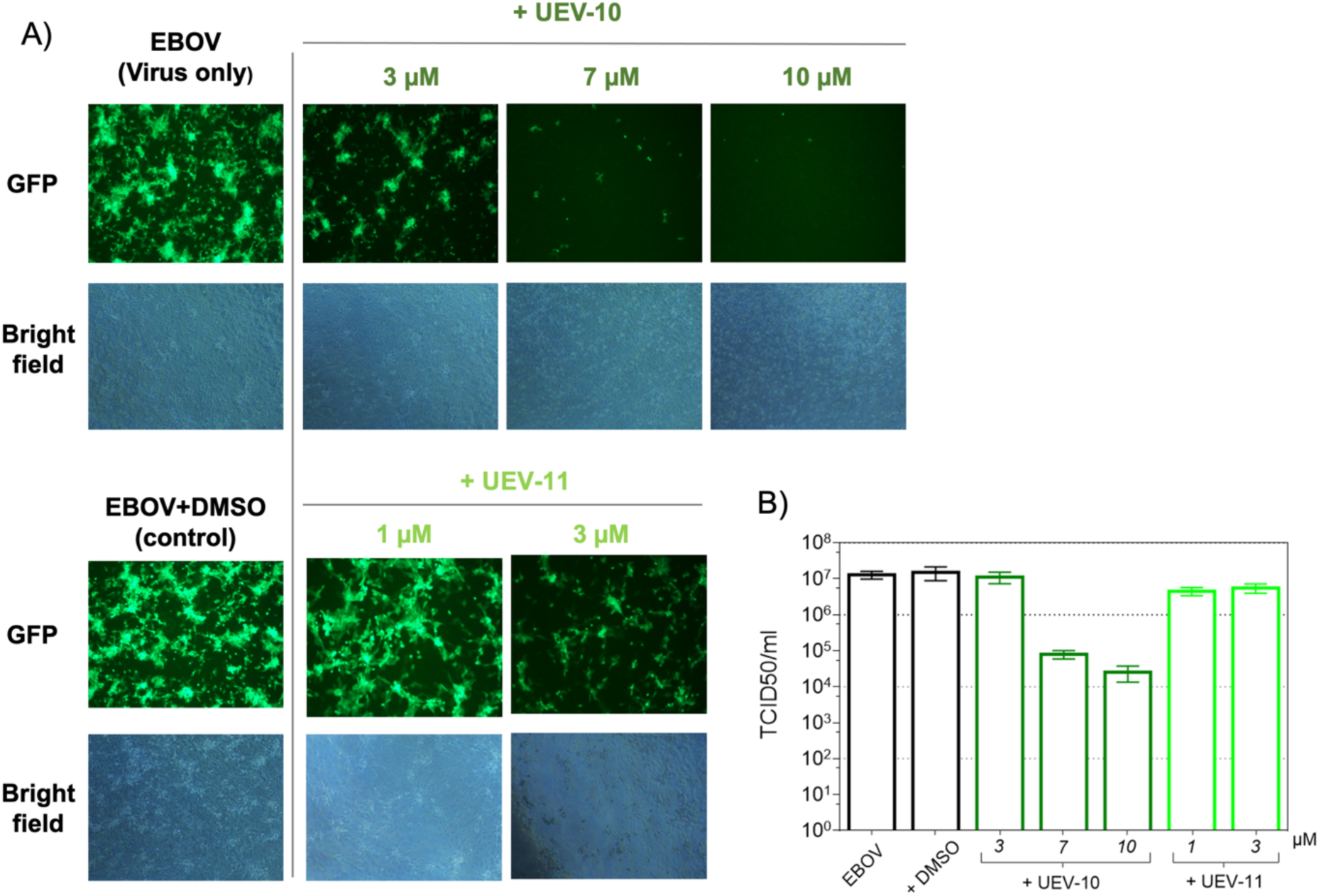
Inhibition of EBOV infection. Vero E6 cells were infected with rgEBOV-VP30-GFP GFP (MOI 0.2) in the presence of UEV-10 or UEV-11 in the indicated concentration for 72 h in duplicates. **a)** GFP expression was monitored to visualize virus spread and bright field pictures were taken to monitor virus CPE and presence or absence of toxic effects. Exemplary pictures are presented. b) Supernatants collected were titrated using TCID50, n=2.

To further assess whether the observed antiviral activity is associated to the specific disruption of TSG101/EBOV-VP40 interactions, the ability of the different compounds to impair EBOV budding in mammalian cells was tested using a well-established virus-like particle (VLP) budding assay based on the expression of the EBOV matrix protein VP40 (Liu et al., 2011; Shepley-McTaggart et al., 2020). Budding of EBOV-VP40 VLP was impaired in the presence of UEV-10 and UEV-11 at 1.0 μM with no observed cytotoxicity, as judged by EBOV-VP40 expression levels in cell extracts (Figure 5A). As before, UEV-11 appeared to be toxic to the cells at 10 μM. No significant changes in VLP egress compared to DMSO control were observed for the rest of compounds. These results confirm that UEV-10 and UEV-11 can efficiently block EBOV-VP40-dependent budding in the absence of other elements from the virus (no other viral proteins are expressed in these assays), sustaining the idea that these compounds specifically target ESCRT recruitment by EBOV-VP40.

**Figure 5.**
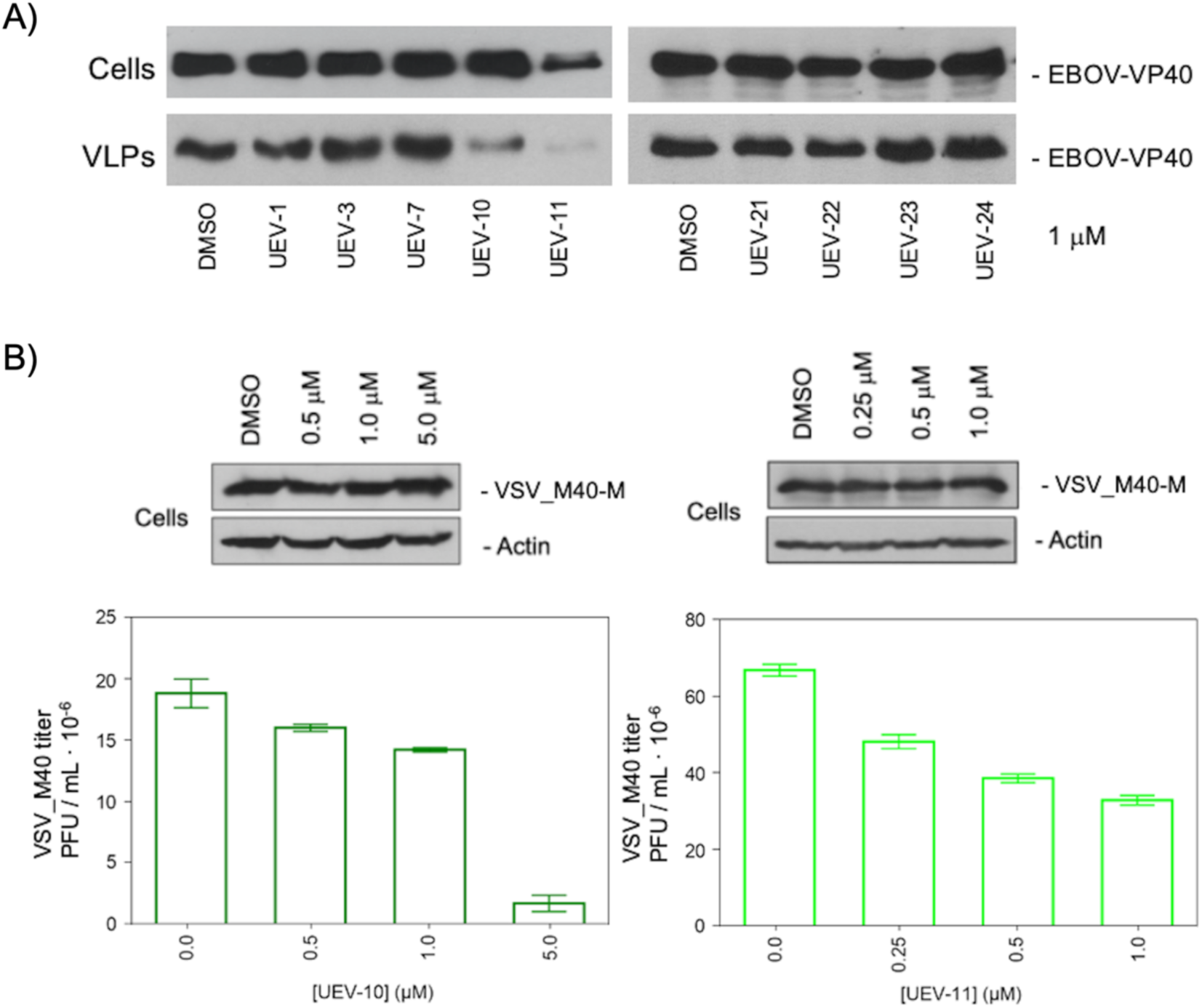
Inhibition of viral budding. **(A)** Virus-like Particle Budding Assays. Various compounds were tested at 1 μM concentration for their ability of inhibit the egress of Ebola EBOV-VP40 virus-like particles. EBOV-VP40 protein was detected in cell extracts and VLPs by Western blot. EBOV-VP40 levels in VLPs were quantified (%) relative to control (DMSO). **(B)** Inhibition of recombinant Vesicular Stomatitis Virus (VSV) budding. HEK293T cells were infected with recombinant virus VSV-M40 at a MOI=0.1 in the presence of DMSO (control) or the indicated concentrations of UEV-10 (left) and UEV-11 (right). Equivalent levels of VSV-M and actin were detected by Western blotting in infected cell extracts (upper panels). Lower panels show the titers of infectious virus released into the supernatants determined on BHK-21 cells (average of 3 independent experiments and the standard error of the mean).

Compounds UEV-10 and UEV-11, showing inhibition of EBOV replication and inhibition of HIV-1 replication in the assay analyzing virus produced RLU, were also tested against a TSG101-dependent Vesicular Stomatitis chimeric virus (VSV-M40) engineered to express the overlapping PPxY and PTAP Late domain sequence from EBOV-VP40 protein (RVIL**PTA<u>P</u>**<u>PEY</u>MEAI) instead of the original PPPY L-domain in its matrix protein (LGIA**PPPY**EEDT) (Irie et al., 2005). As shown in Figure 5B, the two compounds induced significant dose-dependent inhibition of VSV-M40 replication. Western blots of infected cell extracts did not reveal any perturbation on the VSV-M40 and actin levels, indicating that the concentrations of inhibitor used were not cytotoxic and that the observed inhibitory effect was targeted to virus budding without affecting viral or host protein synthesis.

### 2.5. Modelling and docking

Structural models of TSG101-UEV in complex with the different inhibitors were generated using physics-based protein ligand docking methods. Three independent replicas of blind docking calculations were performed taking the TSG101-UEV/EBOV L-domain complex (4EJE) as a reference. The 5 best poses for the different compounds and the corresponding energy values calculated by the Lead Finder and MM-PBSA algorithms are shown in Figure 6 for UEV-10 and in Figures S7-S15 for all compounds. As expected for the docking of small compounds to relatively featureless protein/peptide interaction sites, we find that binding to the TSG101-UEV domain is characterized by a flat energy landscape with shallow energy minima: for all compounds and replicas the energy differences between poses are very small (ΔE_Pose1-Pose2_ average 2.5 ± 0.4 kJ·mol^-1^ for Lead Finder docking and 17 ± 2 kJ·mol^-1^ for MM-PBSA and ΔE_Pose1-Pose5_ average 4 ± 2 kJ·mol^-1^ for Lead Finder docking and 66 ± 6 kJ·mol^-1^ for MM-PBSA). These values are below the thresholds usually accepted as fully discriminatory for these algorithms. In this scenario, additional criteria, such as the establishment of specific interactions known to be critical for ligand recognition in our specific system, must also be considered to select the most probable binding configuration.

**Figure 6.**
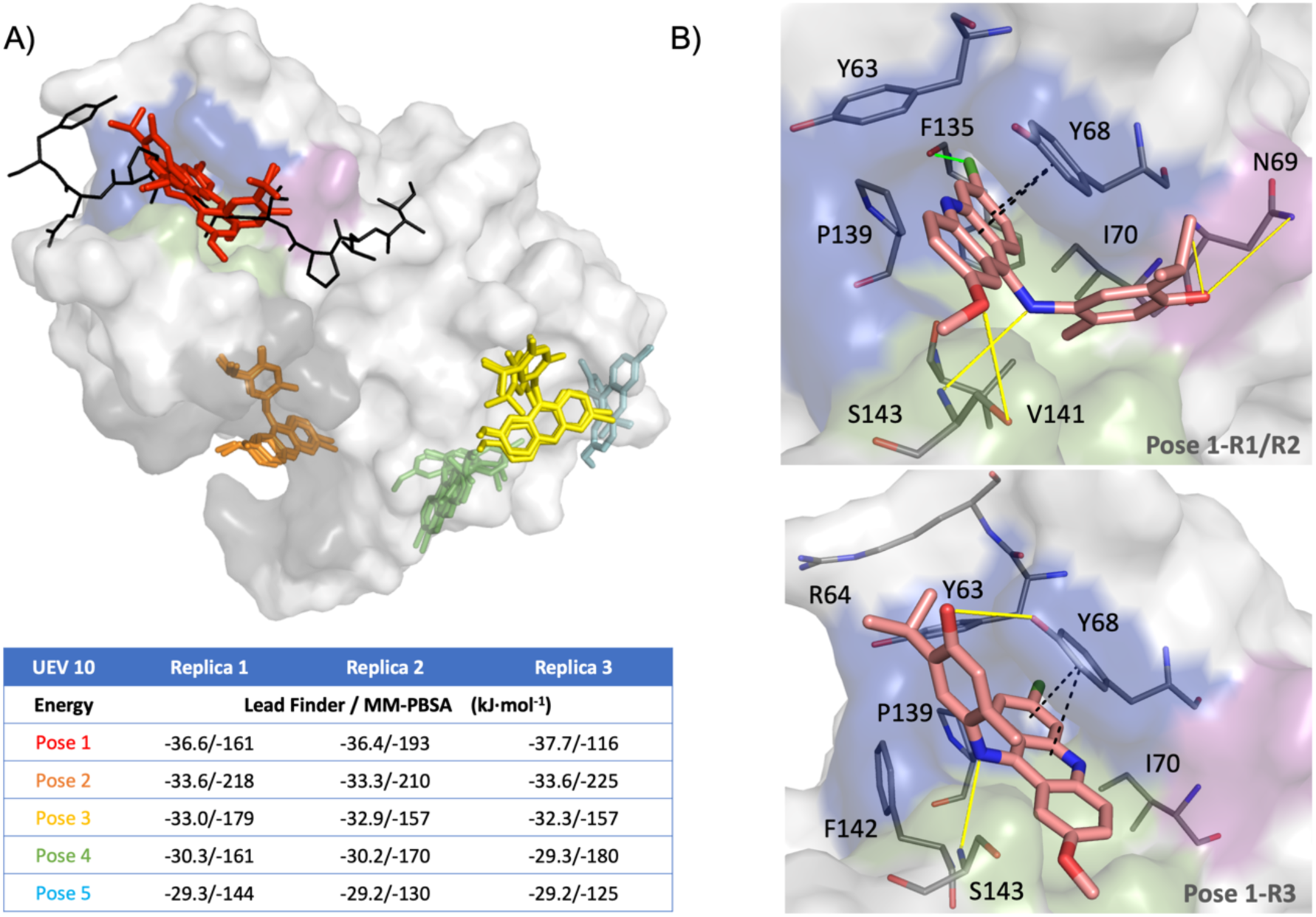
Blind docking of UEV10 compound onto TSG101-UEV. **(A)** Lowest energy poses resulting from three independent replicas of blind docking calculations. Shown is the surface representation of the TSG101-UEV domain (white) in complex with the EBOV L-domain (4EJE.pdb). The PTAP and ubiquitin binding sites are highlighted in color (blue: P0 pocket, green: A-1 pocket, magenta: T-2 pocket; grey: ubiquitin). The top five poses for each replica are shown as colored sticks (Pose 1: red; Pose 2 orange; Pose 3: yellow; Pose 4: green; Pose 5: cyan). The peptide ligand corresponding to the EBOV L-domain in 4EJE is shown in black as a reference. The corresponding energy values for the best 5 poses as calculated by Lead Finder and MM-PBSA for each replica are shown in the Table **(B)** Lowest energy poses for UEV-10 for Replicas 1 & 2 (upper panel) and Replica 3 (lower panel). The most relevant interacting TSG101-UEV residues are shown as grey lines. Protein-ligand interactions calculated by Lead Finder are shown as discontinuous lines (yellow: hydrogen bonds, green: halogen bonds, black: ν-stacking interactions).

Even though the differences in energy are small and non-conclusive, the lowest energy poses generally correspond to conformations compatible with the hypothesis that these compounds target the PTAP site. At this site, all compounds adopt highly reproducible conformations in the three replicas, closely mimicking the interaction pattern characteristic of TSG101-UEV peptide ligands Table 4) (Murciano-Calles et al., 2024). This would be indicative of a competitive inhibition scenario in which the selected compounds disrupt TSG101/virus interaction by displacing TSG101 partners from the PTAP site. The blind docking calculations also produced poses compatible with binding at the ubiquitin site that are within the best 5 poses for all compounds except UEV-1 and UEV-10. Binding at this site would be indicative of an allosteric mechanism of inhibition in which the active compounds would block binding of viral PTAP sequences through long-range cooperative interactions. In most cases, the poses at the ubiquitin site present slightly poorer energies, show more conformational variability between replicas, and present a less defined interaction pattern, typically limited to hydrophobic and hydrogen bonds with a few residues at the “tongue” and “lip” regions of the UEV domain.

**Table 4.**
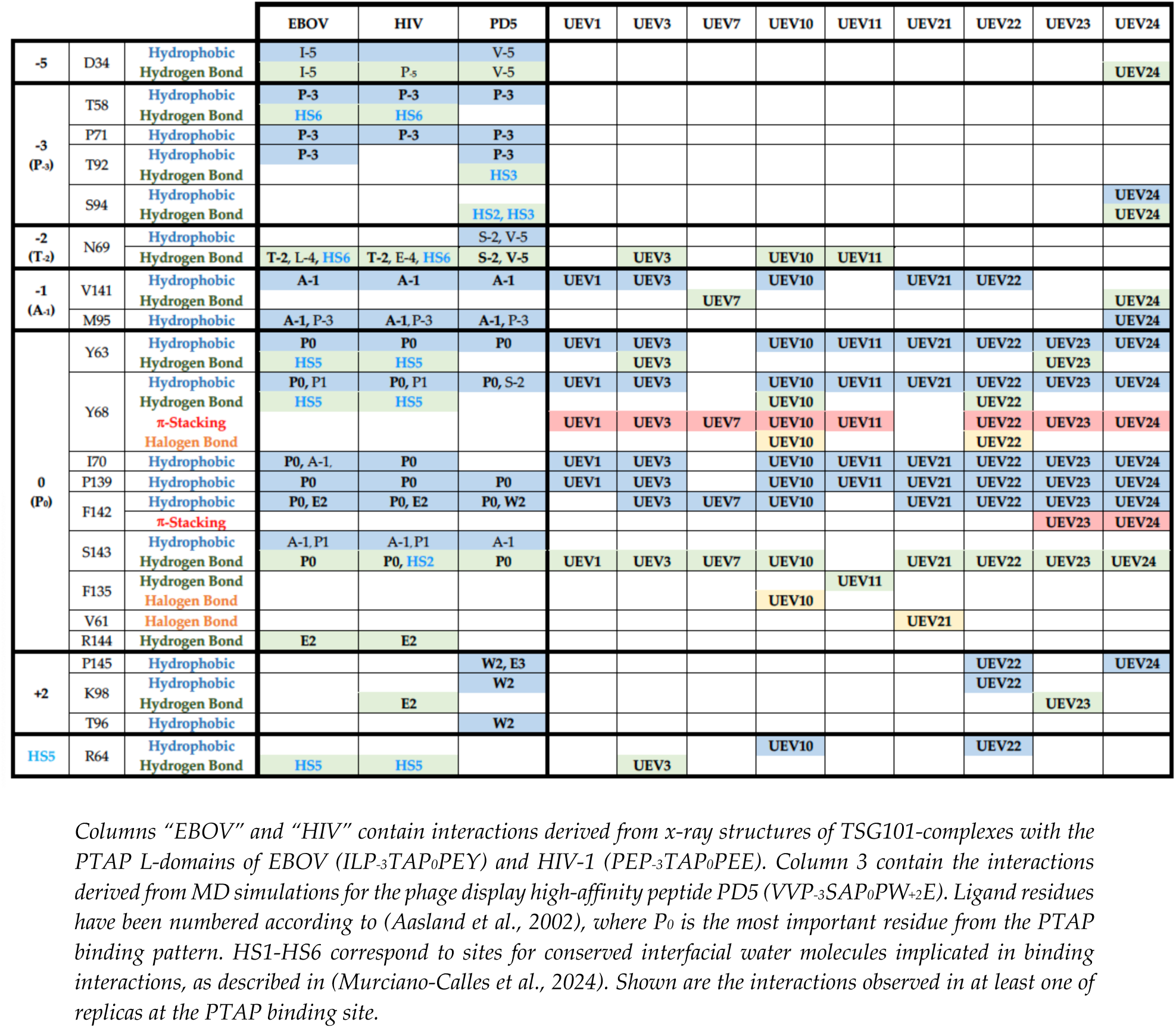
Interactions at the TSG101-UEV PTAP binding site from the lowest energy poses.

At the PTAP site, all compounds occupy a deep hydrophobic pocket formed by residues Y_63_, Y_68_, I_70_, P_139_, F_142_, and S_143_, reminiscent of the “xP pocket” of SH3 and WW domains (Ball et al., 2005).

This pocket is dedicated to the recognition of the second proline within the P_-3_T_-2_A_-1_P_0_ core motif, which is the main anchor point for PTAP recognition. As summarized in Table 5, in all peptide complexes P_0_ establishes strong hydrophobic contacts with residues Y_63_, Y_68_ and F_142_ and a hydrogen bond with S_143_, which are almost invariably reproduced in the PTAP poses for all compounds.

Interestingly, the Y_68_ side chain becomes a key player in small-molecule binding to TSG101-UEV, being systematically implicated in additional interactions not observed in the peptide complexes. These include highly conserved ρε-stacking interactions implicating aromatic rings in the inhibitors that, in the case of the most active compounds (UEV-10 and UEV-22), are complemented by hydrogen and halogen bonds. In addition, the characteristic hydrophobic contacts and hydrogen bonds with V_141_ at the adjacent binding pocket, typically dedicated to A_-1_ recognition, are also frequently observed. The most potent compounds showing antiviral activity (UEV-10, UEV-11, UEV-22, and UEV-24) also reproduce interactions involving residues flanking the PTAP core motif that have been shown to be key for high affinity and specificity for the peptide ligands (Table 4). These include a) hydrogen bonds with the side chain atoms of N_69_ and D_34_, which in the peptide complexes are typically established with residues at positions -4 o -5 at the N-terminal end of the ligand, and b) packing and hydrogen bond interactions with residues and P_145_, K_98_ and T_96_, characteristic of residues at position +2 at the C-terminal region (E_+2_ in HIV-1 and EBOV Late domains and W_+2_ in high affinity phage display ligands). Finally, in some poses, the most active compounds, such as UEV-10, occupy hydration sites that are well defined and highly conserved in the crystal structures of Late domain complexes (Figure 7), establishing hydrogen bond interactions with residues that are typically coordinating interfacial water molecules at hydration sites 2 (S_94_, S_143_) and 5 (Y_63_, Y_68_, R_64_) in the PTAP complex structures (Table 5)(Murciano-Calles et al., 2024).

**Figure 7.**
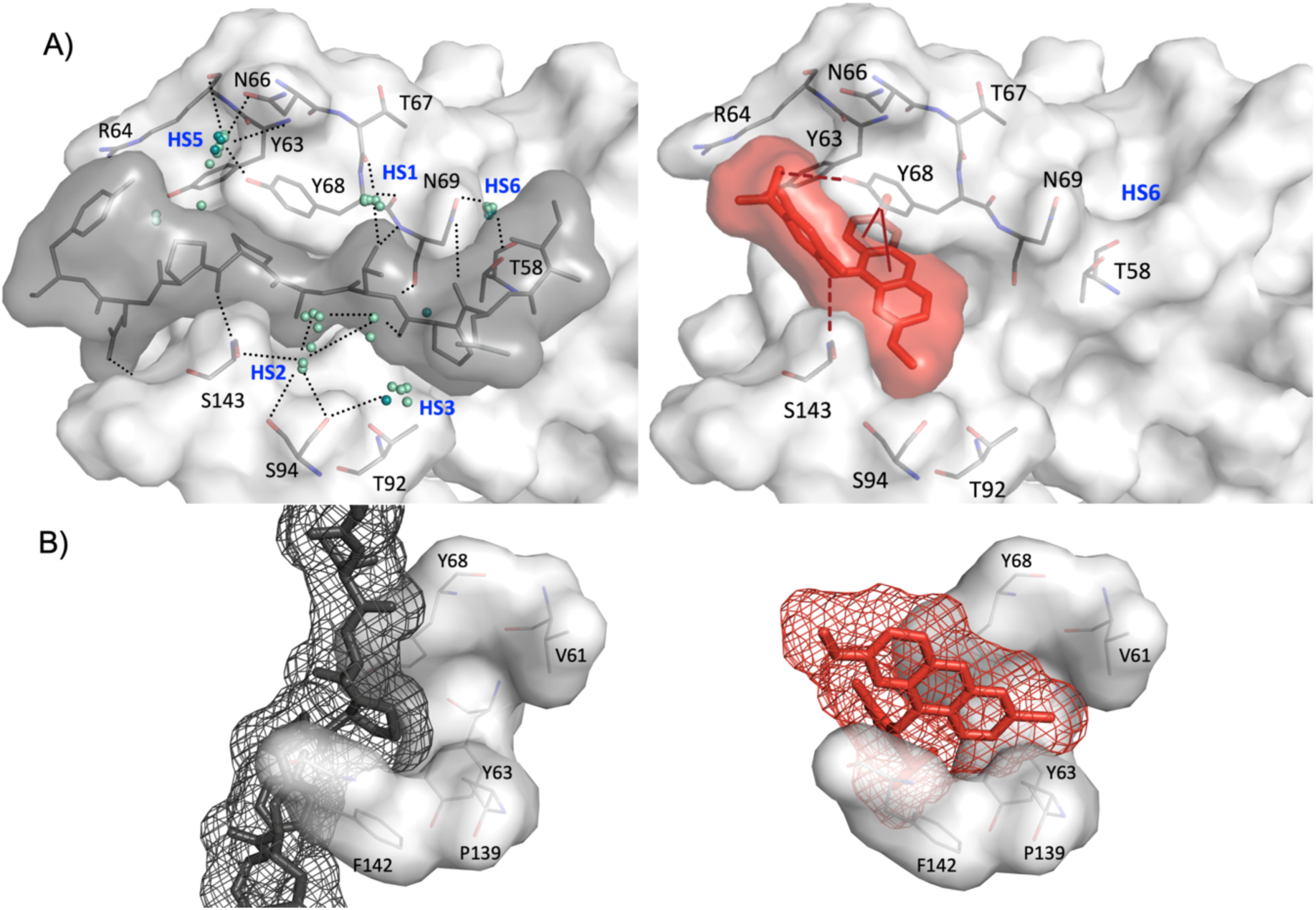
Comparison of the binding modes of PTAP L-domain and small-molecule ligands to TSG101-UEV. **(A)** The left panel shows a surface representation of the TSG101-UEV (white surface) in bound to the EBOV L-domain peptide (grey surface) (4EJE) showing water mediated hydrogen bonds (dotted black lines) established at highly conserved hydration sites (HS1-HS5). The right panel shows the lowest energy pose for compound UEV-10 (red surface) highlighting the most relevant interactions (hydrogen bonds and π-stacking with residues Y_68_ and S_143_) as discontinuous red lines. **(B)** Surface representation of the xP pocket of TSG101-UEV, illustrating how the P_0_ within the PTAP core in the EBOV L-domain (black mesh) remains at the surface while UEV-10 structure (red mesh) inserts deeper into the pocket establishing optimized interactions.

## 3. Discussion

The screening of small-compound libraries against the isolated TSG101-UEV domain using in vitro miniaturized assays has allowed us to identify a set of drug-like ligands that can efficiently block binding of viral L-domain peptides, prevent TSG101 recruitment by the viral proteins in human cells with manageable toxicity, inhibit budding in virus-like particle assays, and, ultimately, impair viral replication.

Notwithstanding the challenge posed by the flat and extended TSG101-UEV/PTAP binding interface, which expands 9-12 amino acid sequences and thousands of Å^2^ (8600 Å^2^ for the EBOV-L-domain), we have identified 4 drug-like scaffolds that bind TSG101-UEV with high affinity and show inhibitory activity in BiMC and antiviral cell-based assays. Despite their small size, these compounds exhibit dissociation constants in the low micromolar range, two orders of magnitude better than the ESCRT-0 HRS cellular partner (Kd = 290 μM) (Im et al., 2010), and 10 times stronger than the HIV-1 (53 μM) and EBOV (23 μM) PTAP L-domain sequences, which, having evolved to displace TSG101/HRS cellular interactions, are the tightest natural UEV ligands known to date. The low μM affinities achieved by the active compounds are in fact comparable to the strongest TSG101-UEV ligands described thus far that include high-affinity phage display peptides (Kd = 1-2 μM) (Murciano-Calles et al., 2024) and peptidomimetics resulting from aromatic functionalization of the first proline in the PTAP motif of the HIV-1 L-domain (Kd = 3.3 μM) (Liu et al., 2008; Kim et al., 2011). These are remarkable results demonstrating that the hot-spot interactions previously identified as being critical for high-affinity binding of PTAP-containing peptides can be efficiently captured by small drug-like molecules.

Only the strongest ligands showed activity in the BiMC assays, suggesting that low micromolar affinity for the UEV domain is required to efficiently block the interaction between full-length TSG101 and the viral proteins containing PTAP L-domains. Also, in agreement with previous reports indicating that viral L-domains function in a context-independent and interchangeable manner (Freed, 2002; Welker et al., 2021), all compounds produced very similar results when tested against the HIV1-Gag polyprotein (a 55 kDa chain containing the 51 aa p6-Gag protein with a PTAP L-domain at its N-terminus and an ALIX-binding LYPX_n_L domain at the C-terminus (Demirov et al., 2002)) or against the EBOV matrix protein VP40 (a 35 kDa oligomeric protein containing three overlapping PPxY, PTAP, and LYPX_n_L L-domains at its N-terminus (Han et al., 2015)). Nonetheless, among the low-micromolar binders, compounds UEV-10 and UEV-11 elicited the strongest dose-dependent effects, while others showed a more modest inhibition. Whether this reflects a less efficient cellular uptake, or a different mechanism of action remains to be elucidated and is out of the scope of this work.

All high affinity inhibitors produced some level of antiviral activity against HIV1 that varied with the cell type and assay used. Nonetheless, only UEV-10 and UEV-11 showed activity against EBOV: both compounds blocked EBOV budding in VLP assays (human HEK293T cells) but only UEV-10 was active in antiviral assays (monkey VERO E6 cells). The origin of this different behavior is unclear. It could be associated to the use of cell lines with different genetic backgrounds (monkey vs human) or with different expression profiles of ESCRT proteins. Nonetheless, it could also reflect differences in the mechanism for ESCRT recruitment and L-domain requirements for the replication of the different viruses. In this way, while the TSG101/PTAP interaction is essential for HIV-1 replication (Garrus et al., 2001), with ALIX playing a secondary role with modest effects on virion production and replication (Fujii et al., 2009)), EBOV L-domains, which target TSG101, ALIX and NEDD4, have been shown to be critical for budding, but not absolutely required for virus replication (L-domain inactivation upon mutation of key residues within the core motif only reduced replication by 1-2 log units (Neumann et al., 2005)). In agreement with this, we find that UEV-10 and UEV-11 can drastically reduce EBOV VLP production at 1μM concentration but higher concentrations (> 7μM for UEV-10) were required to significantly inhibit replication of live EBOV. Interestingly, the UEV-10 dose-response curve tends to saturate at about 2-log reduction in viral titers, paralleling the effect of Late domain inactivation units (Neumann et al., 2005). For HIV-1 and mutant VSV-M40, highly dependent on TSG101 recruitment, significant inhibition was achieved at lower concentrations.

Our small-molecule inhibitors were identified using an experimental set-up devised to select molecules able to disrupt TSG101-UEV/PTAP interactions by binding to the native state of the domain: i.e, compounds stabilizing the protein in the primary DSF screening (ΔTm > 1) that displace HIV1-p6 from the PTAP site in the secondary AlphaScreen assay. Such stabilizing compounds are, by definition, orthosteric inhibitors targeting interaction sites that are binding-competent in the conformation that is predominantly populated in the absence of the ligand. Allosteric inhibitors binding to alternative sites in poorly populated conformations will destabilize the domain resulting in lower Tm values (Velazquez-Campoy et al., 2016). Since a dynamic coupling between the PTAP and ubiquitin sites has been recently established (Strickland et al., 2022), the possibility that the active compounds bound to the ubiquitin site affecting PTAP binding through long-range effects could, in principle, be considered. Nonetheless, this is not a probable scenario within our experimental set-up since it would require changes in the network of intramolecular cooperative interactions and, most likely, the reorganization of the conformational distribution, leading to destabilizing effects. For example, prazole-based TSG101-UEV inhibitors that form covalent adducts at the ubiquitin binding site induced strong destabilizing effects in DSF assays (Strickland et al., 2017; Watanabe et al., 2020).

The chemical nature of the active compounds and the blind docking studies further support interaction at the PTAP site. All families of proline-binding domains rely on the recognition of proline-rich motifs (x**P**xx**P** for SH3 domains, **PP**xY for WW domains, and PTA**P_0_** for UEV domains) containing conserved prolines that cannot be replaced by other natural amino acids without complete loss of binding affinity (P_0_ in the case of UEV domains). This has been traditionally attributed to the cyclic nature of the proline amino acid that reduces the loss in conformational entropy upon binding (Ball et al., 2005). Nonetheless, elegant work by the Lim laboratory demonstrated that proline is selected in vivo not for shape complementarity or rigidity, but due to its unique N-substituted nature that provides specific but low-affinity recognition (Nguyen et al., 1998). Replacement of the conserved prolines by N-substituted moieties that provide better shape complementarity was shown to be an efficient strategy for the design of high affinity inhibitors of SH3 domains (Nguyen et al., 2000). In agreement with these requirements, our best inhibitors contain N-substituted cycles and, according to the docking predictions, are strongly anchored at the xP pocket of TSG101-UEV, where they reproduce the distinctive packing and hydrogen bond pattern observed for residues P_0_ and A_-1_ in the peptide complexes.

Despite being necessary for binding, core motif recognition is not enough to provide high affinity (dissociation constants for the isolated proline-rich motifs typically range 200-300 μM (Iglesias-Bexiga et al., 2019)), and residues flanking the core motif are required for high affinity and specificity. According to the docking results, the active small-molecule inhibitors attain low micromolar binding through different and complementary mechanisms: A) In the first place, they establish optimized interactions at the xP pocket by taking advantage of the unique geometry of the TSG101-UEV xP pocket, which is at the upper region of a deep cylindrical cavity delimited by Y_68_/ F_142_ at the upper level, Y_63_/P_139_ at a middle level and V_61_ at the bottom. While the peptide binding mode forces the pyrrolidine ring of P_0_ to remain packed between Y_68_ and F_142_ at the surface level, our drug-like inhibitors can insert deeper into the pocket (Figure 7B), establishing more extensive packing interactions as well as additional ρε-stacking and hydrogen/halogen bonds with Y_68_ and less frequently with F_142_ that are not attainable for the peptides. Interestingly, previous virtual screening campaigns targeting the TSG-101 PTAP site identified inhibitors containing N-substituted cycles that are probably contributing to their low μM antiviral activity through similar mechanisms (Liu et al., 2011; Lu et al., 2014; Loughran et al., 2016), and significant improvements in binding affinity resulting from the substitution of P_0_ by bulkier N-substituted moieties in peptidomimetic ligands of TSG101-UEV have been reported (Kim et al., 2011). B) Also, some compounds reproduce hot-spot interactions implicating residues flanking the PTAP core, which are required for high affinity and specificity in the peptide complexes. This is observed for the tetracyclic compounds UEV21-24, lacking N-substituted cycles, which mimic the packing and hydrogen bond interactions established by the indole side chain of W_+2_ (strongly selected in phage display experiments) at an alternative pocket adjacent to the PTAP binding site. Interactions at this pocket (T_96_, K_98_, R_144_, P_145_) are essential to achieve low micromolar affinity by the peptide ligands (Murciano-Calles et al., 2024). C) Finally, most compounds displace interfacial water molecules from hydration sites highly conserved in the PTAP complexes. All active compounds strongly interact with the Y_68_ sidechain that, together with Y_63_, N_66_ and R_64_, is implicated in the coordination of a tightly bound water molecule (hydration site 2 in Figure 7A) at the β2-β3 loop in all TSG101-UEV crystal structures, including the free domain and peptide complexes. In some low-energy poses active compounds such as UEV-10 or UEV-22 occupy this pocket, engaging the Y_68_ hydroxyl group in direct hydrogen bonds, and establishing packing interactions with R_64_. Displacement of this tightly coordinated structural water molecule will result in favorable conformational entropy contributions associated to the increase in the degrees of freedom of the released water plus changes in the dynamics of the loops conforming the hydration site. A similar situation is observed for hydration site 2 (residues S_143_, S_94_ and T_92_), located at the base of the A_-1_ pocket implicated in a complex network of water-mediated hydrogen bonds.

In summary, this study provides valuable structural insight into how small-molecule compounds can engage binding hotspots at the PTAP binding site, exploiting the unique features of the TSG101-UEV xP and W_+2_ pockets to unlock its full binding potential, providing excellent opportunities to tune the binding affinity and selectivity through targeted optimization of the chemical and geometric complementarity. In line with previous virtual screening studies suggesting the druggability of the TSG101-UEV/PTAP interface (Liu et al., 2011; Lu et al., 2014), our results demonstrate that simple screening campaigns against the isolated TSG101-UEV domain can lead to the identification drug-like chemical scaffolds with antiviral activity that constitute valuable starting points for the development of novel broad-spectrum antivirals. In fact, our UEV-10 and UEV-11 compounds present structural similarities with drug-repurposing antiviral candidates identified in phenotypic and machine-learning screenings that show antiviral activity against EBOV and SARS-CoV-2 (Ekins et al., 2015; Madrid et al., 2015; Anantpadma et al., 2019; Gorshkov et al., 2021). In this way, quinacrine and pyronaridine (a component of the antimalarial Pyramax) share the UEV-10 acridine scaffold (Lane et al., 2020; Puhl et al., 2022) and the anticancer prodrug lucanthone (Anantpadma et al., 2019) and its metabolized form hycanthone (Gorshkov et al., 2021) share the UEV-11 thioxanthene scaffold. These drug-repurposing compounds also present similarities in their IC50 values (low μM range) and safety profiles: the UEV-10 acridine analogue presents good absorption, metabolism and pharmacokinetic properties (Lane et al., 2019), while thioxanthene analogues of UEV-11 show toxicity above 10 μM (Anantpadma et al., 2019; Gorshkov et al., 2021). Even though the mechanism of action is still not elucidated, these drug-repurposing candidates have been proposed as inhibitors of viral entry due to their strong lysosomotropic character (Pisonero-Vaquero and Medina, 2017). Nonetheless, considering the strong similarities with our UEV-10 and UEV-11 inhibitors, it is reasonable to hypothesize that these compounds might also be acting at later stages of the viral life cycle, inhibiting ESCRT recruitment and viral budding. Since ESCRT (and TSG101) play key roles in membrane repair and organelle homeostasis (Bohannon and Hanson, 2020; Ritter et al., 2022), this is a plausible scenario fully compatible with the lysosomotropic behavior.

In conclusion, the findings of our study have provided insights that will facilitate the development of novel host-oriented broad-spectrum antivirals. A number of chemical scaffolds with inhibitory activity have been selected, and the key hot-spot interactions have been identified, which will be of great value in improving potency and selectivity through targeted optimization. The results presented herein suggest that UEV-10 may serve as a candidate for the hit-to-lead transition in the development of novel host-oriented broad-spectrum antivirals. The compound exhibits low toxicity and displays inhibitory activity against viruses from diverse families, including Retroviridiae, Filoviridiae and Rhabdoviridiae. This activity is observed despite the differing L-domain requirements for ESCRT recruitment, varying replication cycles and distinct pathogenic mechanisms exhibited by these viruses. It is noteworthy that acridine-based UEV-10 analogues demonstrate efficacy against SARS-CoV-2, which is distinguished by a markedly disparate and yet incompletely elucidated budding mechanism. In this regard, the compounds identified herein serve as invaluable instruments to advance our comprehension of ESCRT (or specifically TSG101) cellular functions and its implications in the budding process of disparate viral families.

## 4. Materials and methods

### 4.1. Protein expression and purification

The plasmid pRSETA encoding the TSG101-UEV domain (residues 1–145, Uniprot code Q99816) was generously provided by Dr W. Weissenhorn (EMBL, Grenoble). TSG101-UEV was expressed and purified as described before (Palencia et al., 2006). Briefly, the protein was expressed in Escherichia coli BL21/DE3 codon plus cells as a His-tag fusion protein. 10 mL of an overnight LB preculture was added to 1 L of LB medium supplemented with 50 µg·mL^-1^ ampicillin, 50 µg·mL^-1^ chloramphenicol, and 3 µg·mL^-1^ glucose. The culture was grown at 37 °C to an OD of 0.8. Expression was then induced with 0.2 mM IPTG for 4 h at 37°C, controlling the pH and glucose level (> 1.5 g·L^-1^). Cells were harvested by centrifugation and resuspended in 20 mM sodium phosphate, 500 mM sodium chloride, and 5 mM β-mercaptoethanol at pH 7.4 (RB, resuspension buffer). Cells were frozen in liquid nitrogen, lysed in a French press, and ultracentrifuged at 30000 rpm for 30 min. The supernatant was loaded onto a Ni-NTA column, washed with RB plus 20 mM and 50 mM imidazole, and finally eluted at 500 mM of imidazole. The eluted protein was extensively dialyzed in RB buffer for complete removal of imidazole before a final step of Superdex-75 size-exclusion chromatography (GE-Healthcare, Fairfield, CT). Protein purity was checked by SDS-PAGE and MALDI-TOF mass-spectrometry experiments (Scientific Instrumentation Services of the University of Granada), obtaining a molecular mass of 18215 Da. The extinction coefficient was 24180 cm^-1^·M^-1^ at 280 nm, determined as described elsewhere (Gill and von Hippel, 1989). Purified protein was dialyzed into 50 mM glycine pH 3.0 for flash freezing using liquid nitrogen. Experimental samples were always prepared by extensive dialysis against the corresponding buffers.

The HIV-1 p6 gene was cloned into pGEX-2TK using the EcoRI and BamHI restriction sites and expressed in Escherichia coli BL21 (DE3) cells. LB cultures were incubated at 37 °C. At 0.5-0.7 OD protein expression was induced with 0.3 mM IPTG and the culture was incubated overnight at 20 °C. After harvesting, the cell pellet was suspended in Buffer A (20 mM Tris-HCl, 100 mM NaCl, 10% glycerol, pH 7.4) supplemented with 1% (v/v) Triton X-100, protease inhibitors, and lysozyme (100 µg·mL^-1^). The cells were incubated on ice for 30 min and lysed by sonication. After centrifugation at 13 000 × g for 45 min, the supernatant was filtered using 0.45 µm syringe filters and loaded onto a 1 mL FPLC-GSTrap column (GE Healthcare) pre-equilibrated in Buffer A at a flow rate of 0.5 mL·min^-1^. The column was washed with 30 mL of Buffer A, and the GST-p6 was eluted with 10 mL GST Elution buffer (50 mM Tris-HCl, 150 mM NaCl, 15 mM reduced glutathione, pH 7.4) at a flow rate of 1 mL·min^-1^. The extinction coefficient was 44600 cm^-1^·M^-1^ at 280 nm, determined as previously described (Gill and von Hippel, 1989). The fractions containing the desired protein were combined and dialyzed overnight at 4 °C against Buffer A. Protein aliquots were flash-frozen in liquid nitrogen and stored at −80 °C.

### 4.2. In vitro High Throughput Screening

#### 4.2.1. Library description

The **NCI Diversity Set IV library** is a collection of 1430 natural and synthetic compounds with a diverse structural landscape that have been evaluated as potential anticancer agents. Compounds in the library were selected using the programs Chem-X (Oxford Molecular Group) and Catalyst (Accelrys, Inc.) that use defined pharmacophoric centers (i.e., hydrogen bond acceptor, hydrogen bond donor, positive charge, aromatic, hydrophobic, acid, base) and defined distance intervals to create a finite set of three dimensional, 3-point pharmacophores resulting in over 1,000,000 possible pharmacophores. The selection protocol considered each molecule, all its pharmacophores and each of its conformational isomers. During the generation of the diversity set, the pharmacophores for any candidate compound were compared to the set of all pharmacophores found in structures already accepted into the set. If the current structure had more than 5 new pharmacophores, it was added to the set. An additional objective was to create a diverse set of compounds that were amenable to forming structure-based hypotheses. Thus, molecules that were relatively rigid, with 5 or fewer rotatable bonds, tending to be planar, 1 or less chiral centers, and pharmacologically desirable features (i.e., did not contain: obvious leaving groups, weakly bonded heteroatoms, organometallics, polycyclic aromatic hydrocarbons, etc.) were given priority in the final selection. All compounds were checked for purity via LC/Mass Spec. and found to have a purity of 90% or better by this method. The **NCI Natural Products Set III library** consists of 118 compounds that were selected from the DTP Open Repository collection of 140,000 compounds for structural diversity, containing a variety of scaffold structures having multiple functional groups.

The **ASINEX protein-protein interaction (PPI) library** comprised 7040 molecules of various sizes, frameworks, and shapes ranging from fragment-like entities to macrocyclic derivatives designed as secondary structure mimetics or as epitope mimetics. In the structural mimetics category, they have designed small molecule and macrocyclic scaffolds that mimic the backbone geometry and the projection of side chains as observed in peptide structures at PPI interfaces. The designs cover β-turn/loop mimetics and α-helix mimetics. Since helices present at the interface in 62% of all protein-protein interactions (Bullock et al., 2011), they have focused on designs including mimics with the substitution geometry of an α-helices, as well as designs that mimic the location of “hot-spot” side chains in helix-mediated PPIs. Epitope mimetics were designed using structure-based designs focusing on the locations of hot-spot residues at the PPI interfaces.

The physicochemical properties (molecular weight, partition coefficient LogP, Number of Rotable Bonds, Number of Hydrogen Bond Acceptors and Donors, Topological Polar Surface Area) of the three compound libraries (NCI Diversity Set IV library, Natural Products Set III library and ASINEX PPI library) were calculated using LigandScout v 4.5. The results are detailed in Figure S2.

#### 4.2.2. Differential Scanning Fluorimetry assay

2 µL of the compounds in 100% DMSO at 10 mM concentration were loaded onto 96-well microplates. 23 µL of an assay master mix containing SYPRO Orange and TSG101-UEV in PBS were added to each well to a final assay volume of 25 µL of 800 µM compound, 8% DMSO, 10 µM SYPRO Orange, and 2 µM TSG101-UEV. High affinity peptides PD1 and PD2, previously identified by phage display (Murciano-Calles et al., 2024) were used as positive controls at 800 µM concentration, 8% DMSO. Free TSG101-UEV at 8% DMSO (23 µL of master mix + 2 µL of DMSO) was used as negative control. After 10 min of protein preincubation at RT, unfolding curves were registered from 20 to 95 °C at a 0.5 °C/min scan rate in a Biorad© CFX 96 qPCR real-time thermal cycler using default HEX filter settings. The midpoint unfolding temperature (Tm) was calculated from the first derivative of the curve and compared to the DMSO control to calculate the corresponding Tm shift. Quality parameters for the DSF screening were calculated for each screening plate (Ź-factor ≥ 0.5, signal-to-noise ratio ≥ 9, hit rate ≤ 1%) and primary hits were selected using a ΔTm > 1.0 °C threshold.

#### 4.2.3. AlphaScreen competition assay

An AlphaScreen displacement assay based on the disruption of the His_6_-TSG101-UEV/GST-HIV-p6 interaction (K_d_ = 27 µM) (Garrus et al., 2001) was set up as an orthogonal secondary assay. Purified proteins were extensively dialyzed in PBS supplemented with 0.1% BSA. 1 µM His_6_-TSG101-UEV was pre-incubated with compounds at 800 µM and 8% DMSO for 30 min at RT. GST-HIV-p6 was then added to a final concentration of 0.3 µM and incubated for 30 minutes. Subsequently, nickel chelate acceptor beads and GST conjugate donor beads were added simultaneously to a final concentration of 10 µg·mL^-1^ and incubated for 1 hour at RT in the dark. Assay plates were read in the AlphaScreen mode (excitation wavelength = 680 nm, emission wavelength = 570 nm) on an EnVision plate reader (Revvity). Dose-response experiments for IC_50_ calculations were performed following the same assay protocol (n = 3) and titrating the His6-TSG101-UEV/GST-HIV-p6 interaction against a final compound concentration ranging from 800 to 0.5 µM and at 8% final DMSO concentration along the 14-points dilution series. For IC50 determination, serial dilutions from 800 µM down to 5·10^-4^ µM were prepared in 14 points, maintaining a constant 8% DMSO. IC50 values were obtained from dose-response curves as described elsewhere (Volpe et al., 2014). The Kd of the interaction of the selected inhibitors with TSG101-UEV was determined from the IC50 assuming a competitive binding model, as described elsewhere (Cer et al., 2009) considering a Kd of 27 µM for the His_6_-TSG101-UEV/GST-HIV-p6 interaction (Garrus et al., 2001).

### 4.3. Cytotoxicity assays

Standard MTT tests (Clothier et al., 2013) were performed using HEPG2 cell lines (HB-8065-ATCC). Cells were seeded at 10.000 cells/well in 96-well plates. After 24 h, cells were treated with compounds at final concentrations ranging from 50 to 0.1 μM. MMS (methyl methanesulfonate) 4mM was used as a positive control of 100% cell death and 0.5% DMSO as the corresponding vehicle-alone control. After 72h, 0.5 mg/ml MTT (3-(4,5-dimethylthiazol-2-yl)-2,5-diphenyltetrazolium bromide, Acros Organics, Ref.: #158990050) was added and absorbance at 570 nm was measured after 2 h incubation. Raw data files from the Envision were imported into Genedata Screener® (v21.0.1). Data were normalized using negative and positive controls as reference. Dose-response curves were fitted to a standard Y=Bottom + (Top-Bottom)/(1+10^((X-LogIC50))) model using the Smart Fit function of Screener® and the half-maximal activity derived from the hill equation model (qAC50) as well as the Lower/Upper 95% Confidence Limits (CL) were calculated.

### 4.4. Bimolecular Complementation Assay

The ability of the selected compounds to disrupt the interaction between full-length proteins in a biologically relevant context was assessed using a previously described Bimolecular Complementation Assay (BiMC) (Licata et al., 2003; Liu et al., 2011). The cellular complementation reporter consists in the co-expression of two constructs based on the pCAGGS vector, one encoding the N-terminal domain of YFP (uniprot P42212, residues 1-158) fused to full-length TSG101 (NYFP-TSG101) and the other encoding the C-terminal domain of YFP (residues 159-239) fused to Gag protein from HIV-1 (CYFP-HIV1-Gag) or to the matrix protein VP40 from EBOV (CYFP-eVP40) (Licata et al., 2003). Briefly, HEK293T cells (ECACC, Ref.:#85120602) were seeded overnight at 20,000 cells/well in a BioCoat Poly-D-Lysine Black/Clear Flat Bottom TC-treated 384-well assay plate (Corning®, Ref.:#354663) and then transfected using 5 µL·mL^-1^ lipofectamine LTX (ThermoFisher®, Ref.: #15338100). Cells were co-transfected with a pair of pCAGGS plasmis coding for NYFP-Tsg101 and CYFP-viral protein (CYFP-HIV1-Gag or CYFP-eVP40) constructs at 25 ng/well, incubated for 6 h, and then treated with the selected compounds at a final concentration of 10, 1, 0.5, and 0.1 µM, 0.5% DMSO for 36 h. After treatment, cells were fixed with a 4% formaldehyde solution (Sigma-Aldrich, #Ref.: F1635) in PBS, and their corresponding nuclei were finally stained with 16 µM Hoechst 33342 (Invitrogen, Ref.:#H3570). Stained cells were then examined by an Operetta CLS High Content Analysis System (Revvity®), collecting images with a 20x objective lens, scanning each well in channel 1 for the YFP signal per well (Height = 11 micrometers, Ex = 490 nm, Em = 550 nm, Power = 20%, Exposure = 10 ms, Fields per well = 9) and in channel 2 for the number of viable cells detected by the corresponding Hoechst nuclear staining (Height = 7 micrometers, Ex = 355 nm, Em = 460 nm, Power = 20%, Exposure = 2 ms, Fields per well = 9). Wells containing cells treated with 0.5% DMSO were used as negative controls. All experiments were performed in triplicate in three fully independent replicas, resulting in a total of 9 measurements per each compound/concentration.

### 4.5. Ebola virus VP40 VLP budding assay

HEK293T cells in collagen-coated six-well plates were transfected with plasmids encoding for Ebola virus VP40 (EBOV-VP40) using Lipofectamine reagent (Invitrogen). At 6 h post-transfection, the culture media were replaced with OPTI-MEM in the presence of the indicated compounds and cells were incubated for 24 h. Culture media was harvested, and particles were centrifuged at 351 x g for 10 min to remove cellular debris, and then layered onto a 20% sucrose cushion in STE buffer and centrifuged at 220,000 x g for 2 h at 4°C. The EBOV-VP40 VLP pellet was suspended in STE buffer at 4°C overnight. Cell extracts were prepared using RIPA buffer (50mM Tris HCl pH 8, 150 mM NaCl, 1% NP-40, 0.5% sodium deoxycholate, 0.1% SDS) containing protease inhibitors). The EBOV-VP40 protein in cell lysates and VLPs was analyzed by Western blot with Fisher anti-EBOV-VP40 rabbit polyclonal antibody (primary) and Goat-anti-rabbit HRP (secondary). EBOV-VP40 protein in VLPs was quantified using NIH Image-J software.

### 4.6. Cell-based antiviral assays against recombinant Vesicular Stomatitis Virus (VSV-M40)

The inhibitory activity of the selected compounds on the release of VSV-M40 virus was assessed as described before (Irie et al., 2005). VSV-M40 is chimeric version of Vesicular Stomatitis virus engineered to express the overlapping PPxY and PTAP Late domain sequence from EBOV-VP40 protein (RVIL**PTA<u>P</u>**<u>PEY</u>MEAI) instead of the original PPPY L-domain in its matrix protein (LGIA**PPPY**EEDT) budding through a TSG101-dependen mechanism (Irie et al., 2005). HEK293T cells in collagen-coated six-well plates were infected with VSV-M40 at MOI of 0.1 for 1 h in triplicate. The inoculum was removed, and cells were then washed with three times with PBS. Cells were incubated in OPTI MEM media in the absence or presence of UEV-10 and UEV-11 at the indicated concentrations for 8 h. Virions from the culture media were harvested and centrifuged at 2500 rpm for 10 min at 4°C to remove cellular debris and then titered as described previously (Irie et al., 2005). Cells were lysed in RIPA buffer (50 mM Tris HCl pH 8, 150 mM NaCl, 1% NP-40, 0.5% sodium deoxycholate, 0.1% SDS and protease inhibitor). VSV M protein in cell extracts was detected by SDS-PAGE and Western-blot with anti-VSV M40 monoclonal antibody 23H12, followed by anti-mouse IgG HRP-conjugated secondary antibody both available from Fisher.

### 4.7. Cell-based antiviral assays against HIV-1

#### 4.7.a. Single-cycle infection assays

For these experiments, the conventional TZM-bl neutralization assays were used as previously described (Li et al., 2005). Serial dilutions of peptides were tested on various HIV-1 pseudoviruses and a single cycle infection was detected after 48 h of culture. The inhibitory concentration 50% (IC50) was calculated.

#### 4.7.b. Multiple rounds infection assays

Multiple rounds of HIV replication were measured after 4 days of infection. Briefly, serial dilutions of peptides were incubated with cells and virus and cultured for 4 days. HIV replication was measured with two distinct protocols a) Renilla Luciferase detection following infection of CEM cells or A3R5 cells both expressing CD4 and CCR5 HIV receptors with a competent HIV infectious molecular clones (IMC) of subtype C (Bjox) expressing Renilla luciferase (Sarzotti-Kelsoe et al., 2014). b) detection of infected cells by intracellular measurement of HIV core protein p24 following replication of HIV SF162 virus strain in CEM, A3R5 cell line or Phytohemagglutinin-activated primary Peripheral Blood Mononuclear Cells (PBMC), as previously described (Klingler et al., 2022). Cell toxicity was recorded as described in (Klingler et al., 2022).

### 4.8. Cell-based antiviral assays against Ebola virus

For EBOV infection experiments, Vero E6 cells (kindly provided by Stephan Becker, Philipps-University Marburg) were maintained in Dulbecco’s modified Eagle’s medium (DMEM) supplemented with 100 U/ml penicillin and 1x GlutaMAX and fetal calf serum (10%; DMEM10) for maintenance or 2% for infection experiments (DMEM2) and incubated at 37°C with 5% CO_2_. 2x10^5^ cells per well were seeded and infected with rgEBOV-VP30-GFP (kindly provided by Thomas Hoenen, Friedrich-Loeffler-Institut) (Bodmer and Hoenen, 2022) at a multiplicity of infection (MOI) of 0.2 in the BSL4 laboratory of the Friedrich-Loeffler-Institut (Greifswald-Insel Riems). After 1 h, the inoculum was removed, and cells were washed 3x with 1xPBS. Then, DMEM2 with or without the respective compounds in the indicated concentrations was added. 72 h post infection, GFP expression was monitored using a fluorescent microscope (Nikon ECLIPSE Ts2). Supernatants were collected and stored at -80°C until further use. Virus titers in supernatants were determined by tissue culture infectious dose 50 (TCID50) assay on VeroE6 cells (Wulff et al., 2012).

### 4.9. Modelling and docking

For all compounds selected from the experimental screening, docking simulations were performed using the crystal structure of the TSG101-UEV/EBOV L-domain complex (4EJE), from which ligands, waters and other extraneous molecules had been removed. The modified pdb file was preprocessed using Maestro (Schrödinger Release 2022-1: Desmond Molecular Dynamics System, D. E. Shaw Research, New York, NY, 2021. Maestro-Desmond Interoperability Tools, Schrödinger, New York, NY, 2021. Maestro), utilizing the Protein Preparation Wizard and System builder tools with the OPLS3e forcefield (Roos et al., 2019). The preprocessed structure was saved in mol2 format.

Models of the selected compounds in sdf format were obtained from smi files using Open Babel tool (https://doi.org/10.1186/1758-2946-3-33). Finally, bond orders were assigned and the LigPrep tool from Maestro software was used. The preprocessed molecules were exported to mol2 format.

Blind docking calculations against the protein structure were performed for each compound using the Metascreener in-house tool (https://github.com/bio-hpc/metascreener). The calculations were launched using Lead Finder method (Stroganov et al., 2008). The calculations were performed in CPU nodes from Leftraru supercomputer (NLHPC, https://www.nlhpc.cl/) using 4GB of RAM. Three replicas with different seed were launched in each case. Finally, the complexes obtained from the blind docking calculations for each compound, conformation, and replica were solvated and minimized using ACPYPE (Sousa da Silva and Vranken, 2012) and GROMACS (Abraham et al., 2015). MM/PBSA interaction free energies for the prepared complexes were calculated using the g_mmpbsa software (Kumari et al., 2014).

## 5. Supplementary material description

- **Figure S1:** Concentration and DMSO dependency of the thermal denaturation profiles of TSG101-UEV followed by Differential Scanning Fluorescence.
- **Figure S2:** Physicochemical descriptors of the compound libraries screened in this work.
- **Figure S3:** Cytotoxicity measured by MTT assay for the selected TSG101-UEV/HIV1-p6 inhibitors
- **Figure S4:** Concentration dependence of the disruption of the protein-protein interactions between NYFP-TSG101 and CYFP-EBOV-VP40 or CYFP-HIV1-Gag by compound UEV-10
- **Figure S5:** Concentration dependence of the disruption of the protein-protein interactions between NYFP-TSG101 and CYFP-EBOV-VP40 or CYFP-HIV-1-Gag by compound UEV-11
- **Figure S6:** Effect of compounds on EBOV infection
- **Figure S7:** Blind docking of UEV-1 compound onto TSG101-UEV structure
- **Figure S8:** Blind docking of UEV-3 compound onto TSG101-UEV structure
- **Figure S9:** Blind docking of UEV-7 compound onto TSG101-UEV structure
- **Figure S10:** Blind docking of UEV-10 compound onto TSG101-UEV structure
- **Figure S11:** Blind docking of UEV-11 compound onto TSG101-UEV structure
- **Figure S12:** Blind docking of UEV-21 compound onto TSG101-UEV structure
- **Figure S13:** Blind docking of UEV-22 compound onto TSG101-UEV structure
- **Figure S14:** Blind docking of UEV-23 compound onto TSG101-UEV structure
- **Figure S15:** Blind docking of UEV-24 compound onto TSG101-UEV structure
- **Table S1:** Identity and structure of the compounds selected from the HTS showing activity as inhibitors of TSG101-UEV/HIV1-p6 interactions
- **Table S2**: SwissADME prediction of physicochemical properties, drug-likeness, and medicinal chemistry for the selected TSG101-UEV/HIV1-p6 inhibitors
- **Table S3**.- Prediction of ADME properties for TSG101-UEV/HIV1-p6 inhibitors

## 6. Acknowledgements

This work was supported by the MCIN/AEI/10.13039/501100011033/ and FEDER Una manera de hacer Europa [grant numbers BIO2016-78746-C2-1-R, PID2020-112895RB-I00]; the FEDER/Junta de Andalucía-Consejería de Economía y Conocimiento [grant number CV20-19149]; and the Consejería de Universidad, Investigación e Innovación and FEDER Andalusia Program 2021-2027 [grant number C-EXP-295-UGR23].

Supercomputing resources in this work have been supported by the Plataforma Andaluza de Bioinformática of the University of Málaga, by the supercomputing infrastructure of the NLHPC (ECM-02, Powered@NLHPC), and by the Extremadura Research Centre for Advanced Technologies (CETA−CIEMAT), funded by the European Regional Development Fund (ERDF). CETA−CIEMAT belongs to CIEMAT and the Government of Spain. We thank the support of the C.I.C. of the University of Granada. F. M was supported by a FPU fellowship from the Spanish Ministry of education. J.M.C. acknowledges a reincorporation research contract from the University of Granada.

Authors thank Dr. Hoenen for providing the rgEBOV-VP30-GFP for this study and Dr. Ali Tavassoli for his collaboration and scientific advice on TSG101 assays.

## References

1. Vietri M, Radulovic M, Stenmark H (2020) The many functions of escrts. Nat Rev Mol Cell Biol 21:25–42. PMID: 31705132 [Medline]

2. Calistri A, Reale A, Palu G, Parolin C (2021) Why cells and viruses cannot survive without an escrt. Cells 10. PMID: 33668191 [Medline]

3. Lemus L, Goder V (2022) Membrane trafficking: Escrts act here, there, and everywhere. Curr Biol 32:R292–R294. PMID: 35349820 [Medline]

4. Lu Q, Hope LW, Brasch M, Reinhard C, Cohen SN (2003) Tsg101 interaction with hrs mediates endosomal trafficking and receptor down-regulation. Proc Natl Acad Sci U S A 100:7626–7631. PMID: 12802020 [Medline]

5. Meng B, Lever AML (2021) The interplay between escrt and viral factors in the enveloped virus life cycle. Viruses 13. PMID: 33672541 [Medline]

6. Rivera-Cuevas Y, Carruthers VB (2023) The multifaceted interactions between pathogens and host escrt machinery. PLoS Pathog 19:e1011344. PMID: 37141275 [Medline]

7. Ahmed I, Akram Z, Iqbal HMN, Munn AL (2019) The regulation of endosomal sorting complex required for transport and accessory proteins in multivesicular body sorting and enveloped viral budding - an overview. Int J Biol Macromol 127:1–11. PMID: 30615963 [Medline]

8. Ferraiuolo RM, Manthey KC, Stanton MJ, Triplett AA, Wagner KU (2020) The multifaceted roles of the tumor susceptibility gene 101 (tsg101) in normal development and disease. Cancers (Basel) 12. PMID: 32075127 [Medline]

9. Rose KM (2021) When in need of an escrt: The nature of virus assembly sites suggests mechanistic parallels between nuclear virus egress and retroviral budding. Viruses 13. PMID: 34199191 [Medline]

10. Freed EO (2002) Viral late domains. J Virol 76:4679–4687. PMID: 11967285 [Medline]

11. Bieniasz PD (2006) Late budding domains and host proteins in enveloped virus release. Virology 344:55–63. PMID: 16364736 [Medline]

12. Welker L, Paillart JC, Bernacchi S (2021) Importance of viral late domains in budding and release of enveloped rna viruses. Viruses-Basel 13. PMID: WOS:000689931200001 [Medline]

13. Tavassoli A, Lu Q, Gam J, Pan H, Benkovic SJ, Cohen SN (2008) Inhibition of hiv budding by a genetically selected cyclic peptide targeting the gag-tsg101 interaction. ACS chemical biology 3:757–764. PMID: 19053244 [Medline]

14. Han ZY, Lu JH, Liu YL, Davis B, Lee MS, Olson MA, Ruthel G, Freedman BD, Schnell MJ, Wrobel JE, Reitz AB, Harty RN (2014) Small-molecule probes targeting the viral ppxy-host nedd4 interface block egress of a broad range of rna viruses. J Virol 88:7294–7306. PMID: WOS:000337240700017 [Medline]

15. Loughran HM, Han Z, Wrobel JE, Decker SE, Ruthel G, Freedman BD, Harty RN, Reitz AB (2016) Quinoxaline-based inhibitors of ebola and marburg vp40 egress. Bioorg Med Chem Lett 26:3429–3435. PMID: 27377328 [Medline]

16. Lennard KR, Gardner RM, Doigneaux C, Castillo F, Tavassoli A (2019) Development of a cyclic peptide inhibitor of the p6/uev protein–protein interaction. ACS chemical biology 14:1874–1878.

17. Han Z, Ye H, Liang J, Shepley-McTaggart A, Wrobel JE, Reitz AB, Whigham A, Kavelish KN, Saporito MS, Freedman BD, Shtanko O, Harty RN (2021) Compound fc-10696 inhibits egress of marburg virus. Antimicrob Agents Chemother 65:e0008621. PMID: 33846137 [Medline]

18. Castillo F, Corbi-Verge C, Murciano-Calles J, Candel AM, Han Z, Iglesias-Bexiga M, Ruiz-Sanz J, Kim PM, Harty RN, Martinez JC, Luque I (2022) Phage display identification of nanomolar ligands for human nedd4-ww3: Energetic and dynamic implications for the development of broad-spectrum antivirals. Int J Biol Macromol 207:308–323. PMID: 35257734 [Medline]

19. Pornillos O, Alam SL, Davis DR, Sundquist WI (2002a) Structure of the tsg101 uev domain in complex with the ptap motif of the hiv-1 p6 protein. Nat Struct Biol 9:812–817. PMID: 12379843 [Medline]

20. Pornillos O, Alam SL, Rich RL, Myszka DG, Davis DR, Sundquist WI (2002b) Structure and functional interactions of the tsg101 uev domain. EMBO J 21:2397–2406.

21. Palencia A, Martinez JC, Mateo PL, Luque I, Camara-Artigas A (2006) Structure of human tsg101 uev domain. Acta Crystallogr D Biol Crystallogr 62:458–464. PMID: 16552148 [Medline]

22. Garrus JE, von Schwedler UK, Pornillos OW, Morham SG, Zavitz KH, Wang HE, Wettstein DA, Stray KM, Côté M, Rich RL, Myszka DG, Sundquist WI (2001) Tsg101 and the vacuolar protein sorting pathway are essential for hiv-1 budding. Cell 107:55–65.

23. Katzmann DJ, Babst M, Emr SD (2001) Ubiquitin-dependent sorting into the multivesicular body pathway requires the function of a conserved endosomal protein sorting complex, escrt-i. Cell 106:145–155. PMID: 11511343 [Medline]

24. Sundquist WI, Schubert HL, Kelly BN, Hill GC, Holton JM, Hill CP (2004) Ubiquitin recognition by the human tsg101 protein. Molecular Cell 13:783–789.

25. Strickland M, Watanabe S, Bonn SM, Camara CM, Starich MR, Fushman D, Carter CA, Tjandra N (2022) Tsg101/escrt-i recruitment regulated by the dual binding modes of k63-linked diubiquitin. Structure 30:289–299 e286. PMID: 35120596 [Medline]

26. Ferreon JC, Hilser VJ (2003) Ligand-induced changes in dynamics in the rt loop of the c-terminal sh3 domain of sem-5 indicate cooperative conformational coupling. Protein Sci 12:982–996. PMID: 12717021 [Medline]

27. Palencia A, Camara-Artigas A, Pisabarro MT, Martinez JC, Luque I (2010) Role of interfacial water molecules in proline-rich ligand recognition by the src homology 3 domain of abl. J Biol Chem 285:2823–2833. PMID: 19906645 [Medline]

28. Martin-Garcia JM, Ruiz-Sanz J, Luque I (2012) Interfacial water molecules in sh3 interactions: A revised paradigm for polyproline recognition. Biochem J 442:443–451. PMID: 22115123 [Medline]

29. Iglesias-Bexiga M, Palencia A, Corbi-Verge C, Martin-Malpartida P, Blanco FJ, Macias MJ, Cobos ES, Luque I (2019) Binding site plasticity in viral ppxy late domain recognition by the third ww domain of human nedd4. Sci Rep 9:15076. PMID: 31636332 [Medline]

30. Im YJ, Kuo L, Ren X, Burgos PV, Zhao XZ, Liu F, Burke TR, Jr., Bonifacino JS, Freed EO, Hurley JH (2010) Crystallographic and functional analysis of the escrt-i /hiv-1 gag ptap interaction. Structure 18:1536–1547. PMID: 21070952 [Medline]

31. Murciano-Calles J, Rodriguez-Martinez A, Palencia A, Andujar-Sanchez M, Iglesias-Bexiga M, Corbi-Verge C, Buzon P, Ruiz-Sanz J, Martinez JC, Perez-Sanchez H, Camara-Artigas A, Luque I (2024) Phage display identification of high-affinity ligands for human tsg101-uev: A structural and thermodynamic study of ptap recognition. Int J Biol Macromol 274:133233. PMID: 38901510 [Medline]

32. Aasland R, Abrams C, Ampe C, Ball LJ, Bedford MT, Cesareni G, Gimona M, Hurley JH, Jarchau T, Lehto VP, Lemmon MA, Linding R, Mayer BJ, Nagai M, Sudol M, Walter U, Winder SJ (2002) Normalization of nomenclature for peptide motifs as ligands of modular protein domains. FEBS Lett 513:141–144. PMID: 11911894 [Medline]

33. Ball LJ, Kuhne R, Schneider-Mergener J, Oschkinat H (2005) Recognition of proline-rich motifs by protein-protein-interaction domains. Angew Chem Int Ed Engl 44:2852–2869. PMID: 15880548 [Medline]

34. Liu F, Stephen AG, Waheed AA, Aman MJ, Freed EO, Fisher RJ, Burke TR, Jr. (2008) Sar by oxime-containing peptide libraries: Application to tsg101 ligand optimization. Chembiochem 9:2000–2004. PMID: 18655064 [Medline]

35. Kim SE, Liu F, Im YJ, Stephen AG, Fivash MJ, Waheed AA, Freed EO, Fisher RJ, Hurley JH, Burke TR, Jr. (2011) Elucidation of new binding interactions with the tumor susceptibility gene 101 (tsg101) protein using modified hiv-1 gag-p6 derived peptide ligands. ACS Med Chem Lett 2:337–341. PMID: 21643473 [Medline]

36. Liu Y, Lee MS, Olson MA, Harty RN (2011) Bimolecular complementation to visualize filovirus vp40-host complexes in live mammalian cells: Toward the identification of budding inhibitors. Adv Virol 2011. PMID: 22102845 [Medline]

37. Lu J, Han Z, Liu Y, Liu W, Lee MS, Olson MA, Ruthel G, Freedman BD, Harty RN (2014) A host-oriented inhibitor of junin argentine hemorrhagic fever virus egress. J Virol 88:4736–4743. PMID: 24522922 [Medline]

38. Ertl P, Rohde B, Selzer P (2000) Fast calculation of molecular polar surface area as a sum of fragment-based contributions and its application to the prediction of drug transport properties. J Med Chem 43:3714–3717. PMID: WOS:000089725000014 [Medline]

39. Lipinski CA, Lombardo F, Dominy BW, Feeney PJ (2001) Experimental and computational approaches to estimate solubility and permeability in drug discovery and development settings. Adv Drug Deliv Rev 46:3–26. PMID: 11259830 [Medline]

40. Baell JB, Holloway GA (2010) New substructure filters for removal of pan assay interference compounds (pains) from screening libraries and for their exclusion in bioassays. J Med Chem 53:2719–2740. PMID: WOS:000276096300004 [Medline]

41. Brenk R, Schipani A, James D, Krasowski A, Gilbert IH, Frearson J, Wyatt PG (2008) Lessons learnt from assembling screening libraries for drug discovery for neglected diseases. Chemmedchem 3:435–444. PMID: WOS:000254264200010 [Medline]

42. Sarzotti-Kelsoe M, Daniell X, Todd CA, Bilska M, Martelli A, LaBranche C, Perez LG, Ochsenbauer C, Kappes JC, Rountree W, Denny TN, Montefiori DC (2014) Optimization and validation of a neutralizing antibody assay for hiv-1 in a3r5 cells. J Immunol Methods 409:147–160. PMID: 24607608 [Medline]

43. Klingler J, Paul N, Laumond G, Schmidt S, Mayr LM, Decoville T, Lambotte O, Autran B, Bahram S, Moog C, Cohort AC (2022) Distinct antibody profiles in hla-b *57+, hla-b *57-hiv controllers and chronic progressors. AIDS 36:487–499. PMID: 34581307 [Medline]

44. Shepley-McTaggart A, Fan H, Sudol M, Harty RN (2020) Viruses go modular. J Biol Chem 295:4604–4616. PMID: 32111739 [Medline]

45. Irie T, Licata JM, Harty RN (2005) Functional characterization of ebola virus l-domains using vsv recombinants. Virology 336:291–298. PMID: 15892969 [Medline]

46. Demirov DG, Orenstein JM, Freed EO (2002) The late domain of human immunodeficiency virus type 1 p6 promotes virus release in a cell type-dependent manner. J Virol 76:105–117. PMID: 11739676 [Medline]

47. Han Z, Madara JJ, Liu Y, Liu W, Ruthel G, Freedman BD, Harty RN (2015) Alix rescues budding of a double ptap/ppey l-domain deletion mutant of ebola vp40: A role for alix in ebola virus egress. J Infect Dis 212 Suppl 2:S138–145. PMID: 25786915 [Medline]

48. Fujii K, Munshi UM, Ablan SD, Demirov DG, Soheilian F, Nagashima K, Stephen AG, Fisher RJ, Freed EO (2009) Functional role of alix in hiv-1 replication. Virology 391:284–292. PMID: 19596386 [Medline]

49. Neumann G, Ebihara H, Takada A, Noda T, Kobasa D, Jasenosky LD, Watanabe S, Kim JH, Feldmann H, Kawaoka Y (2005) Ebola virus vp40 late domains are not essential for viral replication in cell culture. J Virol 79:10300–10307. PMID: 16051823 [Medline]

50. Velazquez-Campoy A, Sancho J, Abian O, Vega S (2016) Biophysical screening for identifying pharmacological chaperones and inhibitors against conformational and infectious diseases. Curr Drug Targets 17:1492–1505. PMID: 26844568 [Medline]

51. Strickland M, Ehrlich LS, Watanabe S, Khan M, Strub MP, Luan CH, Powell MD, Leis J, Tjandra N, Carter CA (2017) Tsg101 chaperone function revealed by hiv-1 assembly inhibitors. Nat Commun 8:1391. PMID: 29123089 [Medline]

52. Watanabe SM, Ehrlich LS, Strickland M, Li X, Soloveva V, Goff AJ, Stauft CB, Bhaduri-McIntosh S, Tjandra N, Carter C (2020) Selective targeting of virus replication by proton pump inhibitors. Sci Rep 10:4003. PMID: 32132561 [Medline]

53. Nguyen JT, Turck CW, Cohen FE, Zuckermann RN, Lim WA (1998) Exploiting the basis of proline recognition by sh3 and ww domains: Design of n-substituted inhibitors. Science 282:2088–2092. PMID: 9851931 [Medline]

54. Nguyen JT, Porter M, Amoui M, Miller WT, Zuckermann RN, Lim WA (2000) Improving sh3 domain ligand selectivity using a non-natural scaffold. Chem Biol 7:463–473. PMID: 10903934 [Medline]

55. Ekins S, Freundlich JS, Clark AM, Anantpadma M, Davey RA, Madrid P (2015) Machine learning models identify molecules active against the ebola virus in vitro. F1000Res 4:1091. PMID: 26834994 [Medline]

56. Madrid PB, Panchal RG, Warren TK, Shurtleff AC, Endsley AN, Green CE, Kolokoltsov A, Davey R, Manger ID, Gilfillan L, Bavari S, Tanga MJ (2015) Evaluation of ebola virus inhibitors for drug repurposing. ACS Infect Dis 1:317–326. PMID: 27622822 [Medline]

57. Anantpadma M, Lane T, Zorn KM, Lingerfelt MA, Clark AM, Freundlich JS, Davey RA, Madrid PB, Ekins S (2019) Ebola virus bayesian machine learning models enable new in vitro leads. ACS Omega 4:2353–2361. PMID: 30729228 [Medline]

58. Gorshkov K, Chen CZ, Bostwick R, Rasmussen L, Tran BN, Cheng YS, Xu M, Pradhan M, Henderson M, Zhu W, Oh E, Susumu K, Wolak M, Shamim K, Huang W, Hu X, Shen M, Klumpp-Thomas C, Itkin Z, Shinn P, Carlos de la Torre J, Simeonov A, Michael SG, Hall MD, Lo DC, Zheng W (2021) The sars-cov-2 cytopathic effect is blocked by lysosome alkalizing small molecules. ACS Infect Dis 7:1389–1408. PMID: 33346633 [Medline]

59. Lane TR, Dyall J, Mercer L, Goodin C, Foil DH, Zhou H, Postnikova E, Liang JY, Holbrook MR, Madrid PB, Ekins S (2020) Repurposing pyramax(r), quinacrine and tilorone as treatments for ebola virus disease. Antiviral Res 182:104908. PMID: 32798602 [Medline]

60. Puhl AC, Gomes GF, Damasceno S, Godoy AS, Noske GD, Nakamura AM, Gawriljuk VO, Fernandes RS, Monakhova N, Riabova O, Lane TR, Makarov V, Veras FP, Batah SS, Fabro AT, Oliva G, Cunha FQ, Alves-Filho JC, Cunha TM, Ekins S (2022) Pyronaridine protects against sars-cov-2 infection in mouse. ACS Infect Dis 8:1147–1160. PMID: 35609344 [Medline]

61. Lane TR, Massey C, Comer JE, Anantpadma M, Freundlich JS, Davey RA, Madrid PB, Ekins S (2019) Repurposing the antimalarial pyronaridine tetraphosphate to protect against ebola virus infection. PLoS Negl Trop Dis 13:e0007890. PMID: 31751347 [Medline]

62. Pisonero-Vaquero S, Medina DL (2017) Lysosomotropic drugs: Pharmacological tools to study lysosomal function. Curr Drug Metab 18:1147–1158. PMID: 28952432 [Medline]

63. Bohannon KP, Hanson PI (2020) Escrt puts its thumb on the nanoscale: Fixing tiny holes in endolysosomes. Curr Opin Cell Biol 65:122–130. PMID: 32731154 [Medline]

64. Ritter AT, Shtengel G, Xu CS, Weigel A, Hoffman DP, Freeman M, Iyer N, Alivodej N, Ackerman D, Voskoboinik I, Trapani J, Hess HF, Mellman I (2022) Escrt-mediated membrane repair protects tumor-derived cells against t cell attack. Science 376:377–382. PMID: 35446649 [Medline]

65. Gill SC, von Hippel PH (1989) Calculation of protein extinction coefficients from amino acid sequence data. Analytical Biochemistry 182:319–326.

66. Bullock BN, Jochim AL, Arora PS (2011) Assessing helical protein interfaces for inhibitor design. J Am Chem Soc 133:14220–14223. PMID: 21846146 [Medline]

67. Volpe DA, Hamed SS, Zhang LK (2014) Use of different parameters and equations for calculation of ic(5)(0) values in efflux assays: Potential sources of variability in ic(5)(0) determination. AAPS J 16:172–180. PMID: 24338112 [Medline]

68. Cer RZ, Mudunuri U, Stephens R, Lebeda FJ (2009) Ic50-to-ki: A web-based tool for converting ic50 to ki values for inhibitors of enzyme activity and ligand binding. Nucleic Acids Res 37:W441–445. PMID: 19395593 [Medline]

69. Clothier R, Gomez-Lechon MJ, Kinsner-Ovaskainen A, Kopp-Schneider A, O’Connor JE, Prieto P, Stanzel S (2013) Comparative analysis of eight cytotoxicity assays evaluated within the acutetox project. Toxicol In Vitro 27:1347–1356. PMID: 22951948 [Medline]

70. Licata JM, Simpson-Holley M, Wright NT, Han ZY, Paragas J, Harty RN (2003) Overlapping motifs (ptap and ppey) within the ebola virus vp40 protein function independently as late budding domains: Involvement of host proteins tsg101 and vps-4. J Virol 77:1812–1819. PMID: WOS:000180488700017 [Medline]

71. Li M, Gao F, Mascola JR, Stamatatos L, Polonis VR, Koutsoukos M, Voss G, Goepfert P, Gilbert P, Greene KM, Bilska M, Kothe DL, Salazar-Gonzalez JF, Wei X, Decker JM, Hahn BH, Montefiori DC (2005) Human immunodeficiency virus type 1 env clones from acute and early subtype b infections for standardized assessments of vaccine-elicited neutralizing antibodies. J Virol 79:10108–10125. PMID: 16051804 [Medline]

72. Bodmer BS, Hoenen T (2022) Assessment of life cycle modeling systems as prediction tools for a possible attenuation of recombinant ebola viruses. Viruses 14. PMID: 35632785 [Medline]

73. Wulff NH, Tzatzaris M, Young PJ (2012) Monte carlo simulation of the spearman-kaerber tcid50. J Clin Bioinforma 2:5. PMID: 22330733 [Medline]

74. Roos K, Wu C, Damm W, Reboul M, Stevenson JM, Lu C, Dahlgren MK, Mondal S, Chen W, Wang L, Abel R, Friesner RA, Harder ED (2019) Opls3e: Extending force field coverage for drug-like small molecules. J Chem Theory Comput 15:1863–1874. PMID: 30768902 [Medline]

75. Stroganov OV, Novikov FN, Stroylov VS, Kulkov V, Chilov GG (2008) Lead finder: An approach to improve accuracy of protein-ligand docking, binding energy estimation, and virtual screening. J Chem Inf Model 48:2371–2385. PMID: 19007114 [Medline]

76. Sousa da Silva AW, Vranken WF (2012) Acpype - antechamber python parser interface. BMC Res Notes 5:367. PMID: 22824207 [Medline]

77. Abraham MJ, Murtola T, Schulz R, Páll S, Smith JC, Hess B, Lindahl E (2015) Gromacs: High performance molecular simulations through multi-level parallelism from laptops to supercomputers. SoftwareX 1:19–25.

78. Kumari R, Kumar R, Open Source Drug Discovery C, Lynn A (2014) G_mmpbsa--a gromacs tool for high-throughput mm-pbsa calculations. J Chem Inf Model 54:1951–1962. PMID: 24850022 [Medline]

