## Supplementary Materials for "Exploring the druggability of the UEV domain of human TSG101 in search for broad-spectrum antivirals"

1 Department of Physical Chemistry, Institute of Biotechnology, and Unit of Excellence in Chemistry Applied to biomedicine and Environment. School of Sciences, University of Granada, 18071, Granada, Spain.

2 MEDINA Foundation, Avenida del Conocimiento 34, 18016, Granada, Spain.

3 Structural Bioinformatics and High-Performance Computing (BIO-HPC) Research Group, Universidad Católica de Murcia (UCAM), Guadalupe, Spain;

4 Department of Pathobiology, School of Veterinary Medicine, University of Pennsylvania, 3800 Spruce St., Philadelphia, PA 19104, USA.

5 Laboratoire d'ImmunoRhumatologie Moléculaire, UMR_S 1109, Fédération de Médecine Translationnelle de Strasbourg (FMTS), Université de Strasbourg, Strasbourg, France

6 Institute of Novel and Emerging Infectious Diseases, Friedrich-Loeffler Institute, Federal Research Institute of Animal Health, 17493 Greifswald – Insel Riems;

7 Dipartimento di Scienze degli Alimenti e del Farmaco, Università degli Studi di Parma, Viale delle Scienze, 27/A, 43124 Parma, Italy

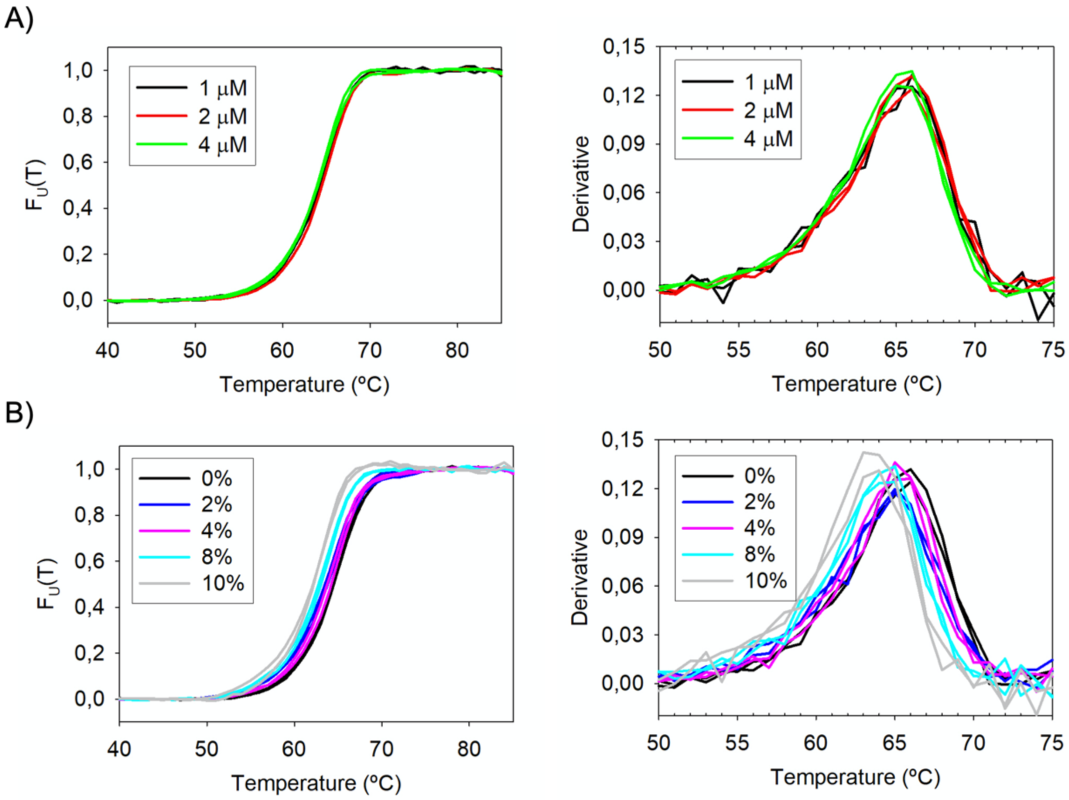


**Figure S1.** Concentration **(A)** and DMSO **(B)** dependency of the thermal denaturation profiles of TSG101-UEV followed by Differential Scanning Fluorescence in PBS buffer, pH 7.5. Left panels show the thermal denaturation profiles at different protein and DMSO concentrations. Right panels show the corresponding first derivative profiles.

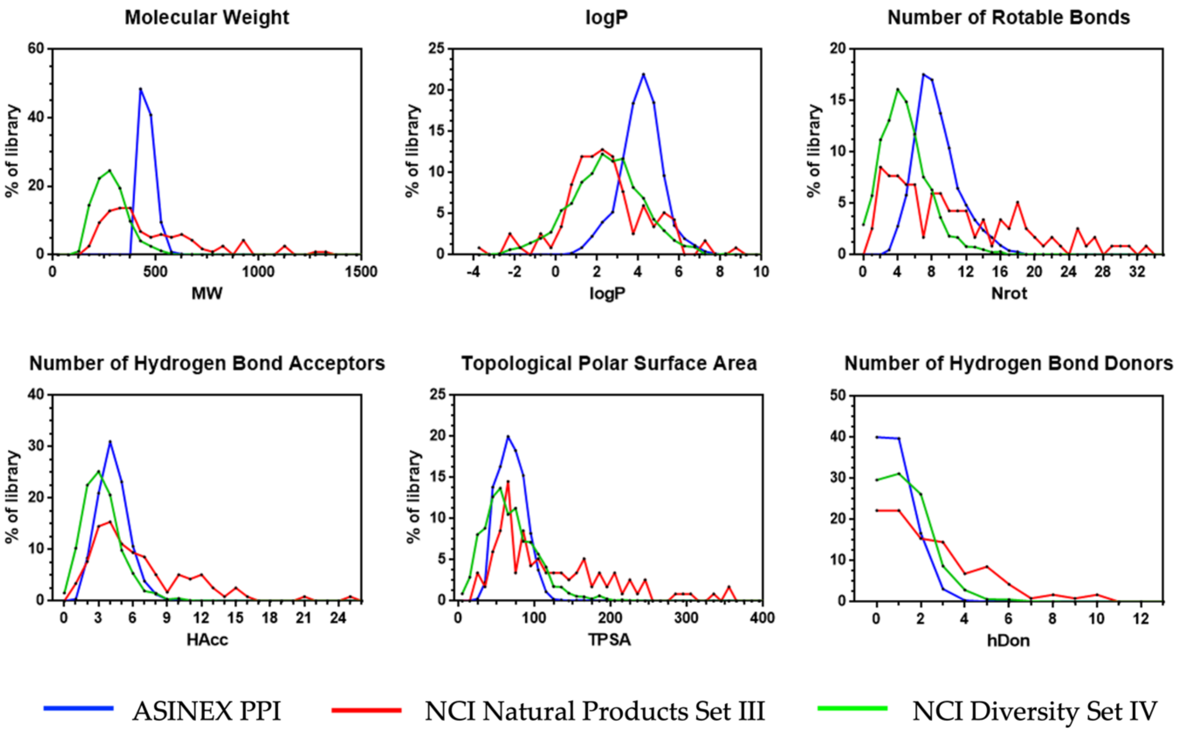


| | NCI Diversity Set IV | NCI Natural Products Set III | ASINEX PP Interaction | All libraries |
|---|---|---|---|---|
| **Number of compounds** | 1596 | 117 | 7040 | 8753 |
| **Molecular Weight (Da)** | 283<br>[169-434] | 482<br>[231-1102] | 458<br>[415-524] | 427<br>[227-521] |
| **Partition Coefficient LogP** | 2.4<br>[-0.6-5.3] | 2.4<br>[-1-5.6] | 4.1<br>[2.2-5.8] | 3.8<br>[1.2-5.7] |
| **Number of rotatable bonds,** | 5.0<br>[1-10] | 10.7<br>[2-27] | 8.6<br>[5-14] | 8<br>[3-14] |
| **Number of H-bond acceptors** | 3.3<br>[1-6] | 6.5<br>[2-15] | 4.2<br>[2-7] | 4.1<br>[2-7] |
| **Number of H-bond donors** | 1.3<br>[0-3] | 2.5<br>[0-8] | 0.8<br>[0-2] | 0.9<br>[0-3] |
| **Topological Polar Surface Area (Å²)** | 68<br>[22-131] | 124<br>[39-300] | 70<br>[42-101] | 70<br>[38-107] |

**Figure S2.-** Physicochemical descriptors of the screened libraries: ASINEX Protein-Protein Interaction (solid blue lines,) NCI Natural Products Set III (solid red lines), and NCI Diversity Set IV (solid green lines) as calculated with LigandScout. The table summarize the average values plus the 5% and 95% percentiles (brackets) for the most relevant physicochemical parameters calculated for the different libraries.

**Table S1.** Compounds selected from the HTS showing activity as inhibitors of TSG101-UEV/HIV1-p6 interactions

| | **UEV-1** | **UEV-3** | **UEV-7** | **UEV-10** | **UEV-11** |
|---|---|---|---|---|---|
| **Library ID** | NSC13974 | NSC369070 | NSC19061 | NSC13051 | NSC317003 |
| **CAS number** | 131-22-6 | NA | 6971-23-9 | 5427-55-4 | 80568-29-2 |
| **Formula** | $C_{16}H_{13}N_3$ | $C_{14}H_9N_3O_3S$ | $C_{18}H_{14}Cl_2N_6$ | $C_{24}H_{23}ClN_2O_2$ | $C_{18}H_{21}IN_2O_2S$ |
| **Structure** | 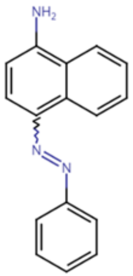 | 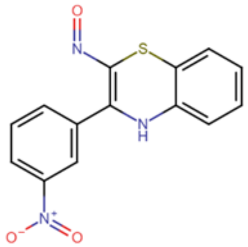 | 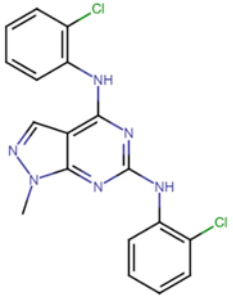 | 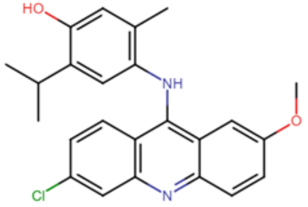 | 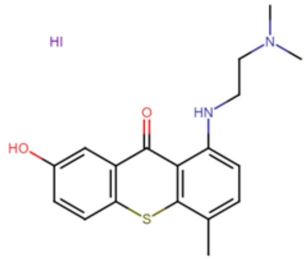 |

| | UEV-21 | UEV-22 | UEV-23 | UEV-24 |
|---|---|---|---|---|
| **Library ID** | BDG33951866 | BDG33905718 | BDG33905712 | BDH33905649 |
| **CAS number** | 2125521-28-8 | 2125521-25-5 | 2127067-05-2 | 2059388-80-4 |
| **Formula** | $C_{32}H_{39}ClN_2O_3$ | $C_{34}H_{43}N_3O_5$ | $C_{34}H_{43}N_3O_6$ | $C_{31}H_{40}N_2O_6$ |
| **Structure** | 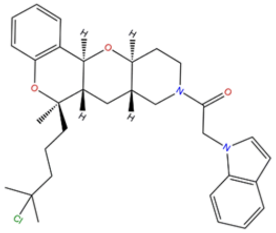 | 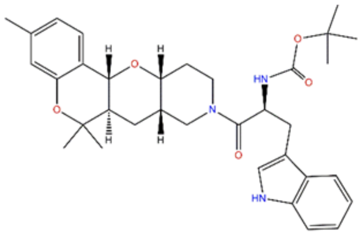 | 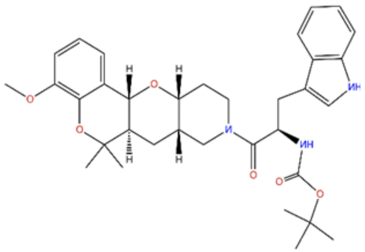 | 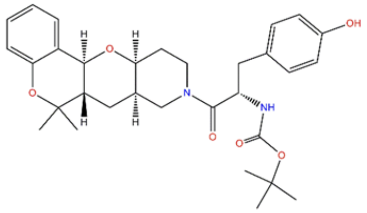 |

**Table S2**.- SwissADME prediction of physicochemical properties, drug-likeness, and medicinal chemistry for the selected TSG101-UEV/HIV1-p6 inhibitors

| | | UEV-1 | UEV-3 | UEV-7 | UEV-10 | UEV-11 |
|---|---|---|---|---|---|---|
| **Physicochemical properties** | Size **MW(Da)** | 247.3 | 299.3 | 385.2 | 406.9 | 456.3 |
| | Polarity **TPSA (Å$^2$)** | 50.74 | 112.58 | 67.66 | 54.38 | 80.81 |
| | Solubility **LogS (ESOL)** | -4.55 | -3.96 | -5.83 | -6.70 | -5.18 |
| | Saturation **Fraction Csp3** | 0.00 | 0.00 | 0.06 | 0.21 | 0.28 |
| | Flexibility **# rotatable bonds** | 2 | 3 | 4 | 4 | 4 |
| | Lipophilicity **LogP (XLOGP3)** | 4.27 | 3.24 | 5.19 | 6.49 | 3.72 |
| **Drug likeness rules** | **Lipinski 1** | 0 | 0 | 1 | 1 | 0 |
| | **Veber 2** | 0 | 0 | 0 | 0 | 0 |
| **Medicinal Chemistry** | **PAINS alerts** | 1 alert: azo_A | -- | -- | 1 alert: amino_acridine_A | -- |
| | **Brenk alerts** | 2 alerts: aniline, diazo_group | 2 alerts: nitro_group, oxygen-nitrogen_single_bond | 0 | 2 alerts: hydroquinone, polycyclic_aromatic_ hydrocarbon_2 | 2 alerts: iodine, polycyclic_aromatic_ hydrocarbon_2 |

| | | UEV-21 | UEV-22 | UEV-23 | UEV-24 |
|---|---|---|---|---|---|
| **Physicochemical properties** | Size<br>**MW(Da)** | 535.1 | 573.7 | 589.7 | 536.7 |
| | Polarity<br>**TPSA (Å²)** | 43.7 | 92.89 | 102.12 | 97.33 |
| | Solubility<br>**LogS (ESOL)** | -6.85 | -6.62 | -6.39 | -5.82 |
| | Saturation<br>**Fraction Csp3** | 0.53 | 0.53 | 0.53 | 0.55 |
| | Flexibility<br>**# rotatable bonds** | 7 | 8 | 9 | 8 |
| | Lipophilicity<br>**LogP (XLOGP3)** | 6.13 | 5.53 | 5.13 | 4.68 |
| **Drug likeness rules** | **Lipinski [1]** | 2 | 1 | 1 | 1 |
| | **Veber [2]** | 0 | 0 | 0 | 0 |
| **Medicinal Chemistry** | **PAINS alerts** | 0 | 0 | 0 | 0 |
| | **Brenk alerts** | 1 alert: alkyl_halide | --- | --- | --- |

**Table S3**.- Prediction of ADME properties for TSG101-UEV/HIV1-p6 inhibitors

| | | UEV1 | UEV3 | UEV7 | UEV10 | UEV11 | UEV21 | UEV22 | UEV23 | UEV24 |
|---|---|---|---|---|---|---|---|---|---|---|
| GI absorption[1] | SwissADME | High | High | High | Low | High | High | High | High | High |
| | admetSAR | High | High | High | High | High | High | High | High | High |
| | pkCSM | High | High | High | High | High | High | High | High | High |
| BBB permeant[2] | SwissADME | Yes | No | No | No | No | No | No | No | No |
| | admetSAR | Yes | Yes | Yes | Yes | Yes | Yes | Yes | Yes | Yes |
| | pkCSM | No | No | No | No | No | No | No | No | No |
| CYP2D6 inhibitor[3] | SwissADME | No | No | Yes | Yes | Yes | Yes | No | No | No |
| | admetSAR | No | No | No | No | No | Yes | No | No | No |
| | pkCSM | No | No | No | Yes | No | No | No | No | No |
| CYP3A4 inhibitor[4] | SwissADME | No | No | Yes | No | Yes | Yes | Yes | Yes | Yes |
| | admetSAR | Yes | Yes | Yes | Yes | No | No | Yes | No | No |
| | pkCSM | No | Yes | Yes | No | Yes | Yes | Yes | Yes | No |
| AMES toxicity[5] | admetSAR | Yes | Yes | Yes | Yes | Yes | No | No | No | No |
| | pkCSM | Yes | No | Yes | Yes | No | No | No | No | No |

[1] *Gastrointestinal absorption;* [2]*Blood-brain barrier permeant;* [3]*Cytochrome P4502D6;* [4]*Cytochrome P45034A;* [5]*Salmonella typhimurium reverse mutation assay*

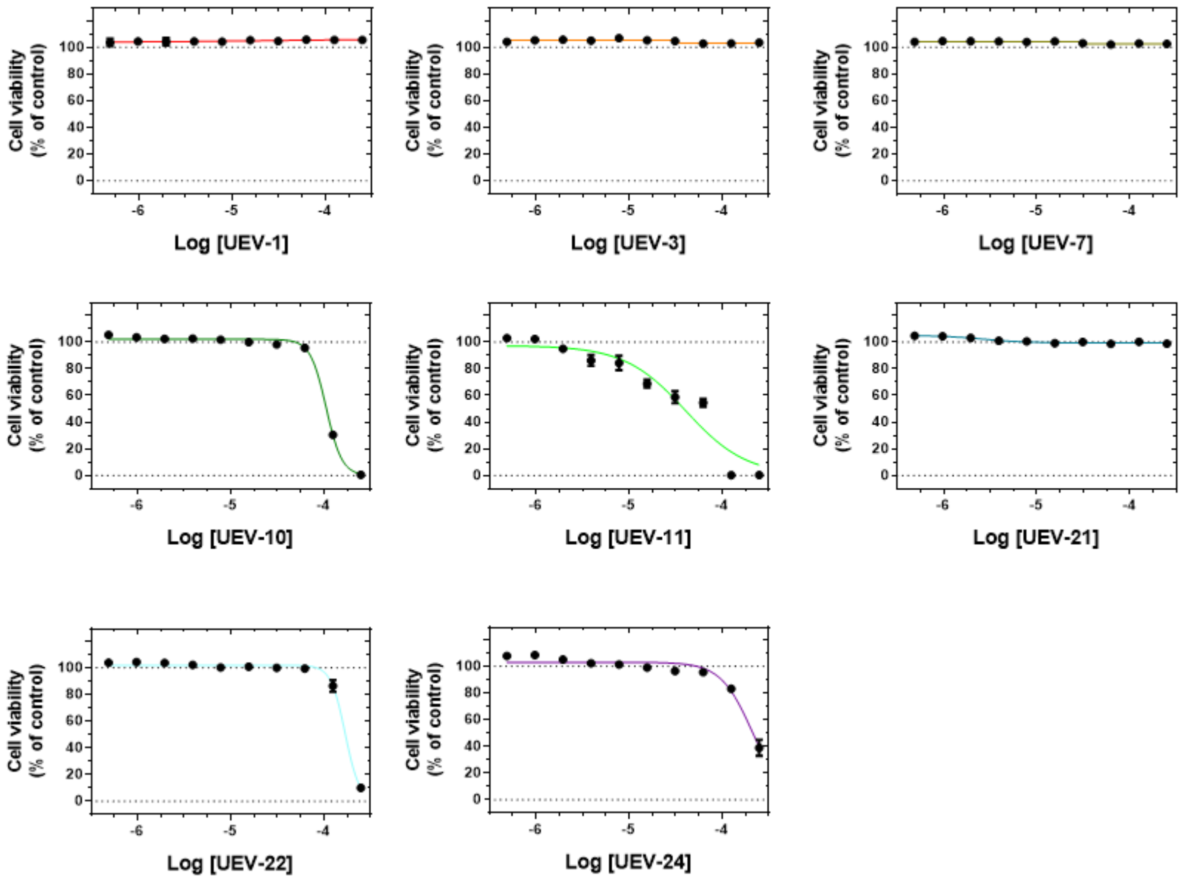


**Figure S3.** Percent of cytotoxicity measured by MTT assay on human HEPG2 cells exposed to the selected TSG101-UEV/HIV1-p6 inhibitors at concentrations ranging from 50 to 0.1 μM upon 2 h incubation. All results were normalized to unexposed controls. Symbols represent the average of three independent experiments and error bars show the standard deviation at a 95% confidence interval. Colored lines represent the best fit to a standard Hill model used to calculate the qAC50 values.

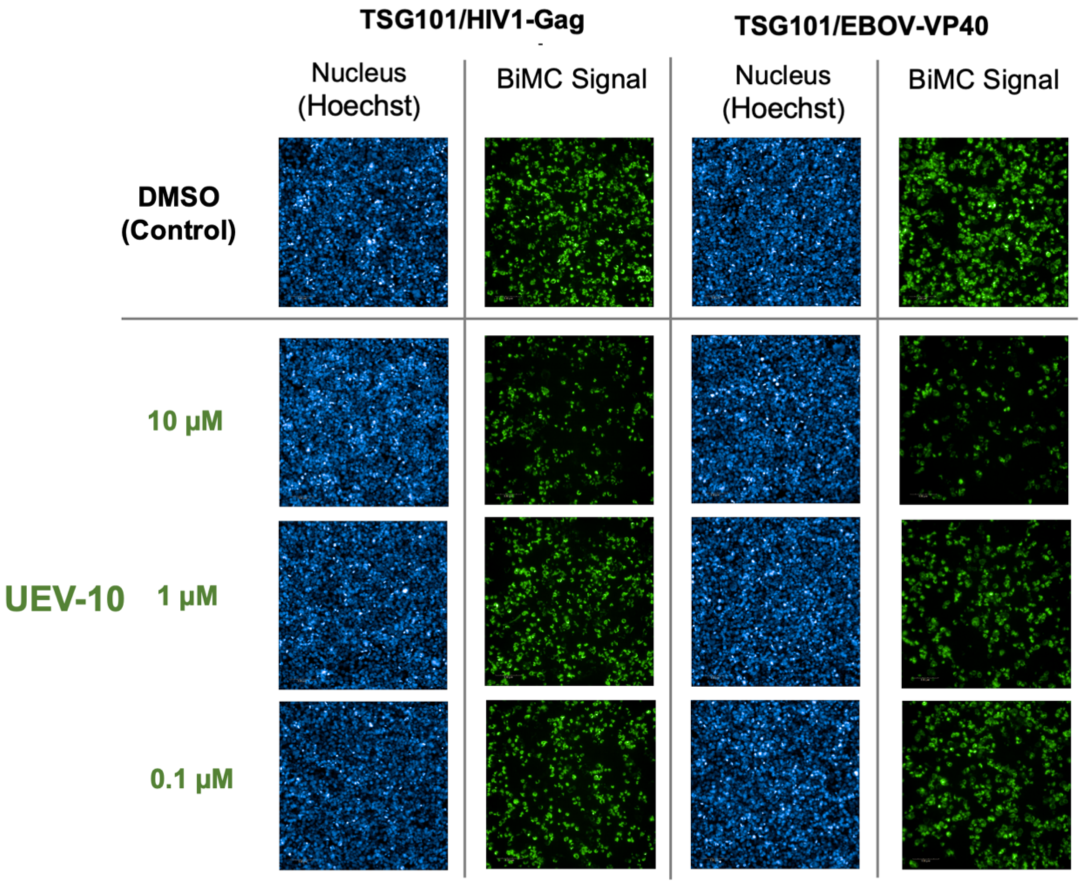


**Figure S4.** Concentration dependence of the disruption of the protein-protein interactions between NYFP-TSG101 and CYFP-EBOV-VP40 or CYFP-HIV1-Gag by compound UEV-10. Shown are representative images of HEK293T cells co-expressing NYFP-TSG101 and CYFP-EBOV-VP40/CYFP-HIV1-Gag fusion proteins treated with UEV-10 compound at different concentrations and 0.5% DMSO. Cells only treated with 0.5% DMSO were taken as negative controls. For each experiment, blue images show the signal corresponding to the Hoechst-stained cell nuclei reporting on the total number of cells in the well and lower green show the fluorescence signal arising from the interaction between NYFP-TSG101 and the viral proteins CYFP-EBOV-VP40 or CYFP-HIV1-Gag.

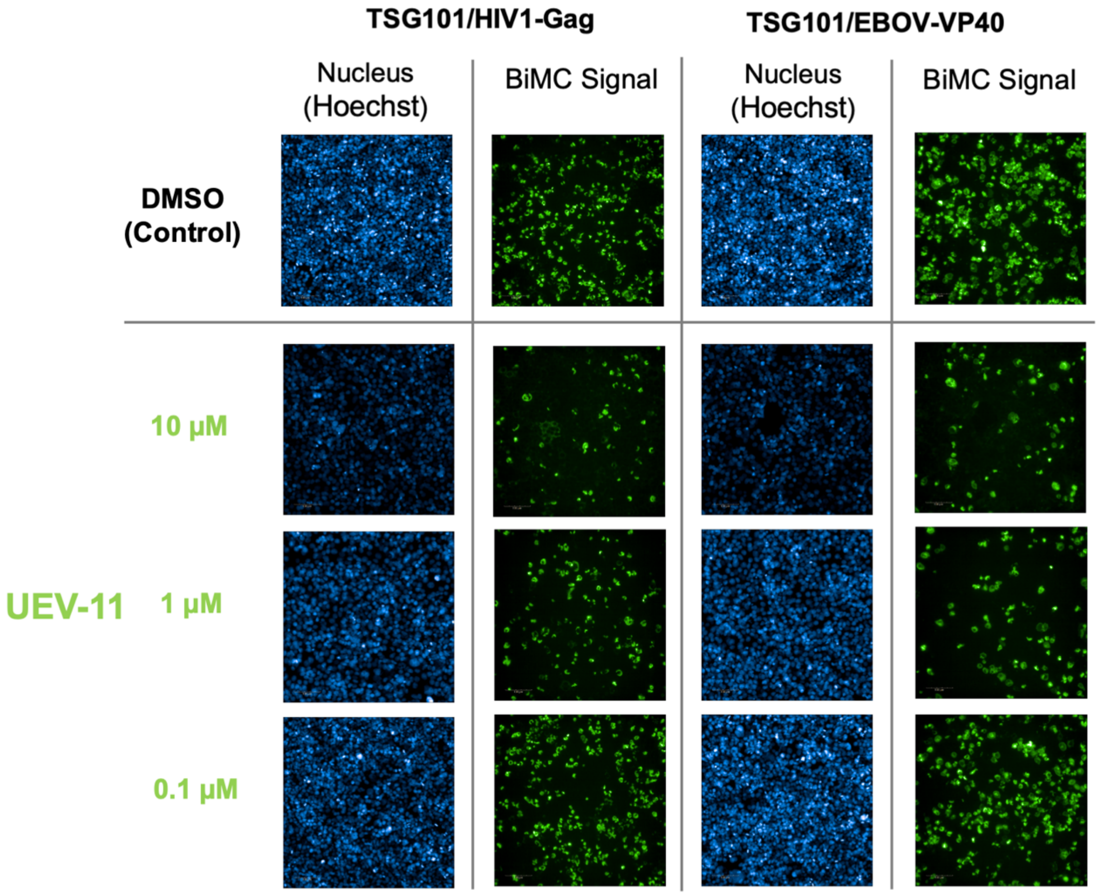


**Figure S5.** Concentration dependence of the disruption of the protein-protein interactions between NYFP-TSG101 and CYFP-EBOV-VP40 or CYFP-HIV-1-Gag by compound UEV-11. Shown are representative images of HEK293T cells co-expressing NYFP-TSG101 and CYFP-EBOV-VP40/CYFP-HIV-1-Gag fusion proteins treated with UEV-11 compound at different concentrations and 0.5% DMSO. Cells only treated with 0.5% DMSO were taken as negative controls. For each experiment, blue images show the signal corresponding to the Hoechst-stained cell nuclei reporting on the total number of cells in the well and lower green show the fluorescence signal arising from the interaction between NYFP-TSG101 and the viral proteins CYFP-EBOV-VP40 or CYFP-HIV-1-Gag.

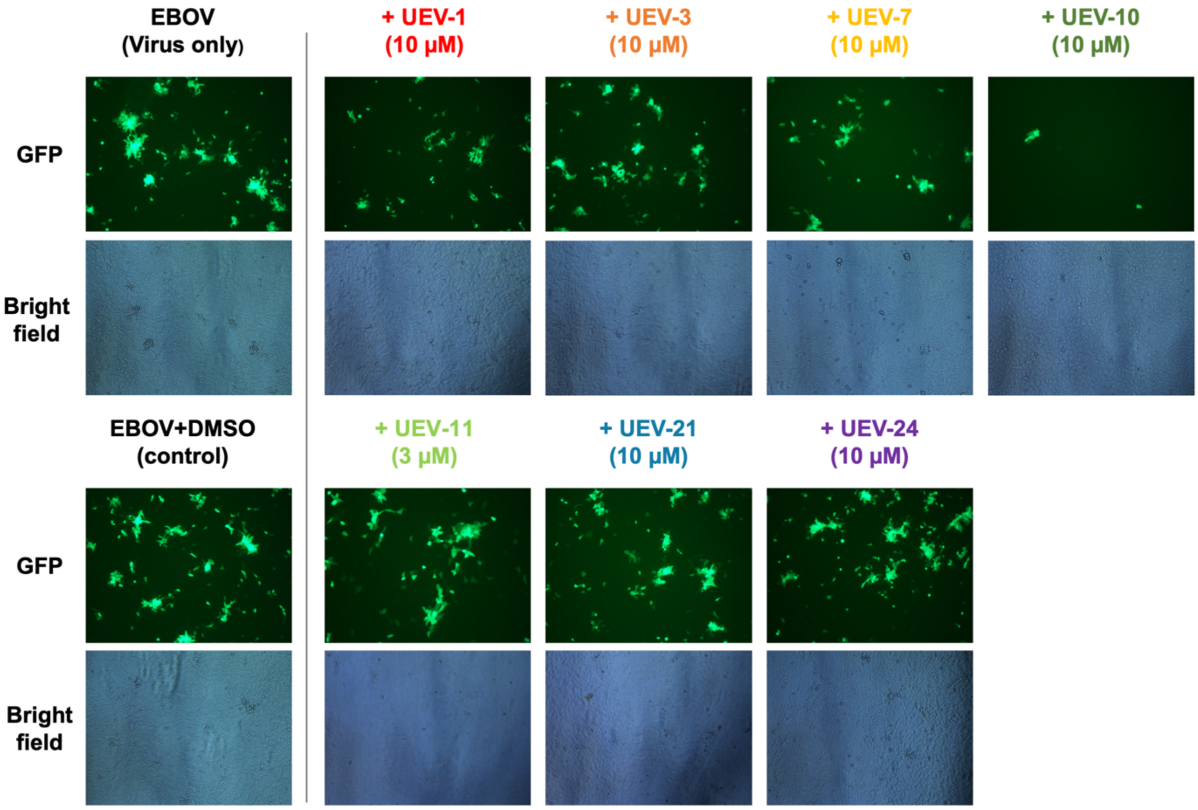


**Figure S6. Effect of compounds on EBOV infection.** Vero E6 cells were infected with rgEBOV-VP30-GFP in the presence of the indicated compounds for 72 h in duplicates. GFP expression was monitored to visualize virus spread. Exemplary pictures are presented.

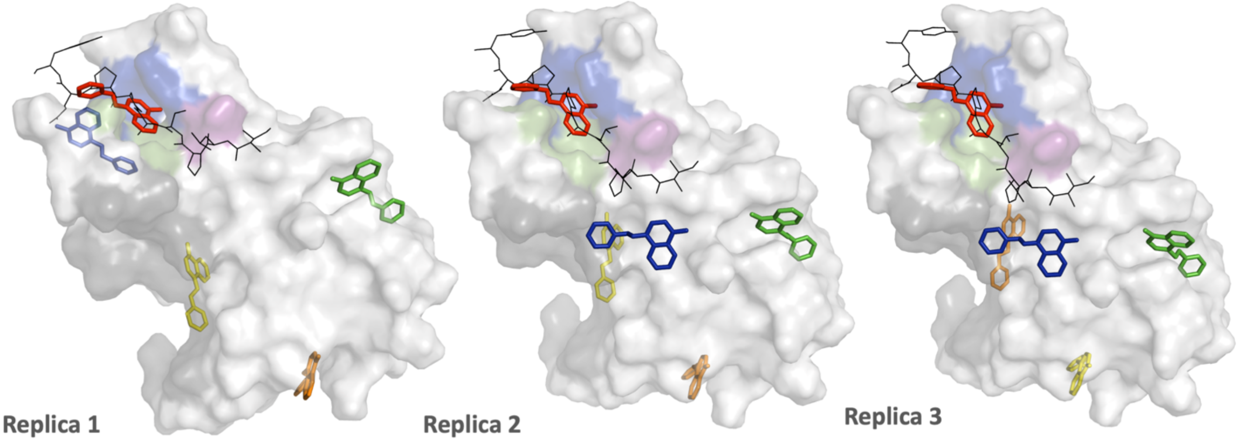


| UEV-1 | Replica 1 | Replica 2 | Replica 3 |
|---|---|---|---|
| Energy | Lead Finder / MM-PBSA (kJ·mol$^{-1}$) | | |
| Pose 1 | -25.6/-141 | -25.6/-143 | -25.7/-136 |
| Pose 2 | -23.1/-148 | -23.2/-146 | -23.3/-133 |
| Pose 3 | -23.0/-131 | -22.9/-149 | -22.8/-129 |
| Pose 4 | -20.4/-158 | -21.8/-137 | -21.8/-142 |
| Pose 5 | -20.3/-123 | -20.3/-146 | -20.0/-120 |

| | PTAP site | Replica 3-Pose 1 | Replica 2-Pose 1 | Replica 1-Pose 1 |
|---|---|---|---|---|
| UEV-1 | Energy LF/MMPBSA (kJ·mol$^{-1}$) | **-25.7 / -136** | **-25.6 / -143** | **-25.5 / -141** |
| | Hydrophobic | Y63, Y68, I70, P139, V141 | Y63, Y68, I70, P139 | Y63, Y68, I70, P139, V141 |
| | Hbonds | S143 | S143 | S143 |
| | π-Stacking | Y68 | Y68 | Y68 |
|  | **Ub site** | **----** | **----** | **---** |

**Figure S7.- Blind docking of UEV-1 compound onto TSG101-UEV.** Shown is the surface representation of the TSG101-UEV domain (white) in complex with the EBOV L-domain (4EJE.pdb). The PTAP and Ubiquitin binding sites are highlighted in color (blue: P0 pocket, green: A-1 pocket, magenta: T-2 pocket; grey: ubiquitin). The top five poses for UEV-1 from each independent replica are shown as colored sticks (Pose 1: red; Pose 2 orange; Pose 3: yellow; Pose 4: green; Pose 5: cyan). The peptide ligand corresponding to the EBOV L-domain in 4EJE is shown in black as a reference. The table summarizes the binding energies calculated by the Lead Finder and MM-PBSA algorithms together with the different forces and interacting residues contributing to binding for the best 3 poses in the PTAP and Ubiquitin sites.

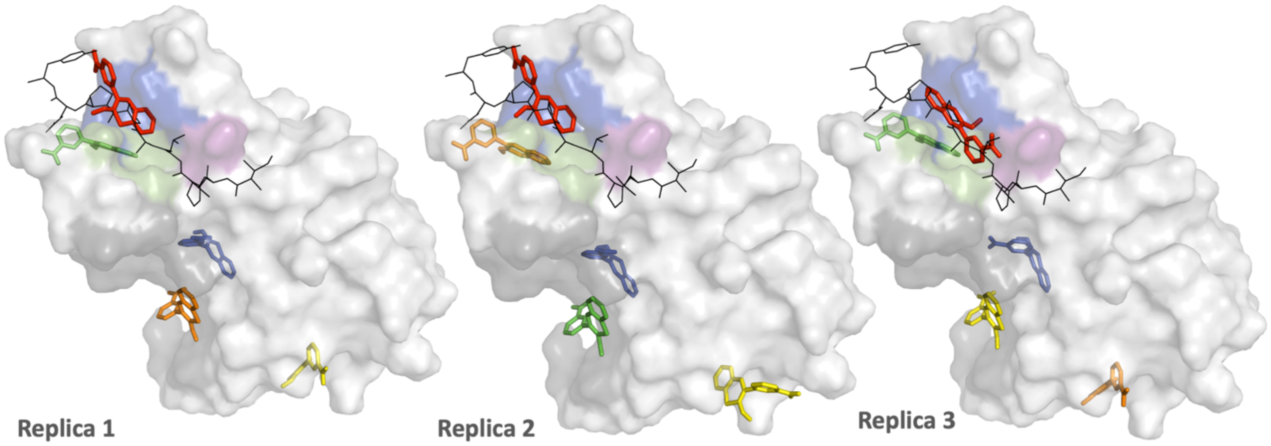


| UEV-3 | Replica 1 | Replica 2 | Replica 3 |
|---|---|---|---|
| **Energy** | **Lead Finder / MM-PBSA (kJ·mol$^{-1}$)** | | |
| Pose 1 | -25.1/-149 | -26.3/-158 | -25.8/-136 |
| Pose 2 | -23.1/-118 | -23.1/-146 | -23.3/-133 |
| Pose 3 | -22.9/-130 | -22.9/-175 | -23.1/-118 |
| Pose 4 | -21.5/-143 | -22.9/-122 | -22.9/-153 |
| Pose 5 | -20.4/-107 | -20.2/-128 | -19.2/-118 |

| | **PTAP site** | **Replica 2-Pose 1** | **Replica 3-Pose 1** | **Replica 1-Pose 1** |
|---|---|---|---|---|
| **UEV-3** | **Energy LF/MMPBSA (kJ·mol$^{-1}$)** | **-26.3 / -158** | **-25.8 / -136** | **-25.1 / -149** |
| | Hydrophobic | Y63, Y68, I70, P139, F142 | Y63, Y68, I70, V141, F142 | Y63, Y68, I70, P139, F142 |
| | Hbonds | Y63, R64, S143 | N69, S143 | Y63, R64, S143 |
| | $\pi$-Stacking | Y68 | | Y68 |
| | **Ub site** | **Replica 3-Pose 3** | **Replica 1-Pose 2** | **Replica 2-Pose 4** |
| | **Energy LF/MMPBSA (kJ·mol$^{-1}$)** | **-23.1 / -118** | **-23.0 / -118** | **-22.9 / -122** |
| | Hydrophobic | **Y42, W75, F88** | **Y42, W75, F88**, K90 | **Y42, W75, F88** |
| | Hbonds | **Y42,** V43, K108 | **Y42**, V43, K108 | **Y42**, V43, K108 |

**Figure S8.- Blind docking of UEV-3 compound onto TSG101-UEV.** Shown is the surface representation of the TSG101-UEV domain (white) in complex with the EBOV L-domain (4EJE.pdb). The PTAP and Ubiquitin binding sites are highlighted in color (blue: P0 pocket, green: A-1 pocket, magenta: T-2 pocket; grey: ubiquitin). The top five poses for UEV-3 from each independent replica are shown as colored sticks (Pose 1: red; Pose 2 orange; Pose 3: yellow; Pose 4: green; Pose 5: cyan). The peptide ligand corresponding to the EBOV L-domain in 4EJE is shown in black as a reference. The table summarizes the binding energies calculated by the Lead Finder and MM-PBSA algorithms together with the different forces and interacting residues contributing to binding for the best 3 poses in the PTAP and Ubiquitin sites.

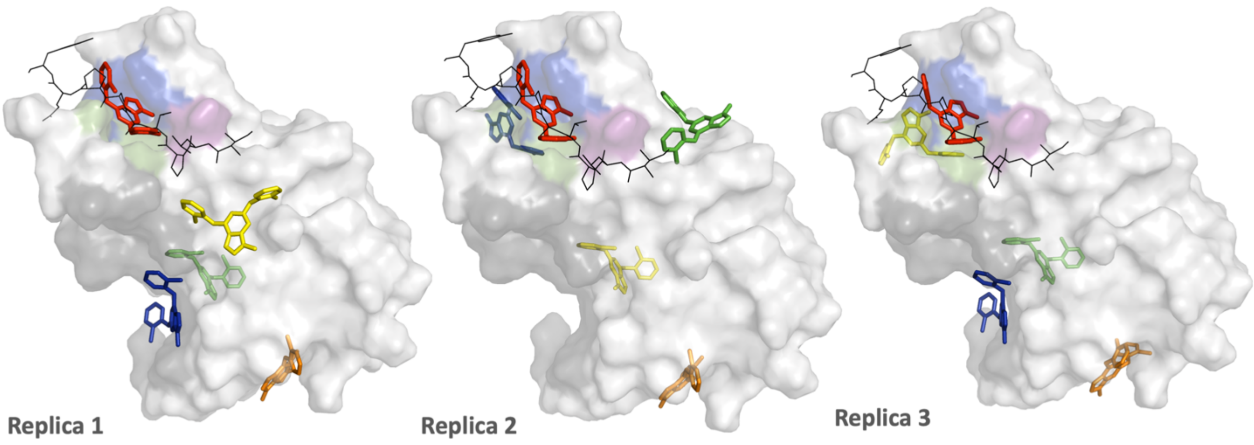


| UEV-7 | Replica 1 | Replica 2 | Replica 3 |
|---|---|---|---|
| Energy | Lead Finder / MM-PBSA (kJ·mol$^{-1}$) | | |
| Pose 1 | -32.8/-169 | -32.9/-164 | -32.7/-202 |
| Pose 2 | -32.4/-171 | -32.1/-168 | -30.3/-167 |
| Pose 3 | -29.3/-129 | -28.7/-132 | -29/-168 |
| Pose 4 | -28.6/-139 | -27.9/-170 | -28.7/-148 |
| Pose 5 | -28.5/-133 | -27.2/-87 | -27.4/-115 |

| | PTAP site | Replica 2-Pose1 | Replica 1-Pose 1 | Replica 3-Pose 1 |
|---|---|---|---|---|
| UEV-7 | **Energy** LF/MMPBSA (kJ·mol$^{-1}$) | **-32.9 / -164** | **-32.7 / -169** | **-32.7 / -202** |
|  | Hydrophobic | F142 | F142 | ---- |
| | Hbonds | V141, S143 | S143 | V141, S143 |
| | π-Stacking | Y68 | Y68 | Y68 |
|  | **Ub site** | **Replica 1-Pose 5** | **Replica 3-Pose 5** | ----- |
| | **Energy** LF/MMPBSA (kJ·mol$^{-1}$) | **-28.5 / -133** | **-27.3 / -115** | |
| | Hydrophobic | **Y42, W75, F88**, K90 | **Y42, W75, F88**, K90 | |
| | Hbonds | V43, **W75** | V43, **W75** | |
| | π-Stacking | **F88** | **F88** | |

**Figure S9.- Blind docking of UEV-7 compound onto TSG101-UEV.** Shown is the surface representation of the TSG101-UEV domain (white) in complex with the EBOV L-domain (4EJE.pdb). The PTAP and Ubiquitin binding sites are highlighted in color (blue: P0 pocket, green: A-1 pocket, magenta: T-2 pocket; grey: ubiquitin). The top five poses for UEV-7 from each independent replica are shown as colored sticks (Pose 1: red; Pose 2 orange; Pose 3: yellow; Pose 4: green; Pose 5: cyan). The peptide ligand corresponding to the EBOV L-domain in 4EJE is shown in black as a reference. The table summarizes the binding energies calculated by the Lead Finder and MM-PBSA algorithms together with the different forces and interacting residues contributing to binding for the best 3 poses in the PTAP and Ubiquitin sites.

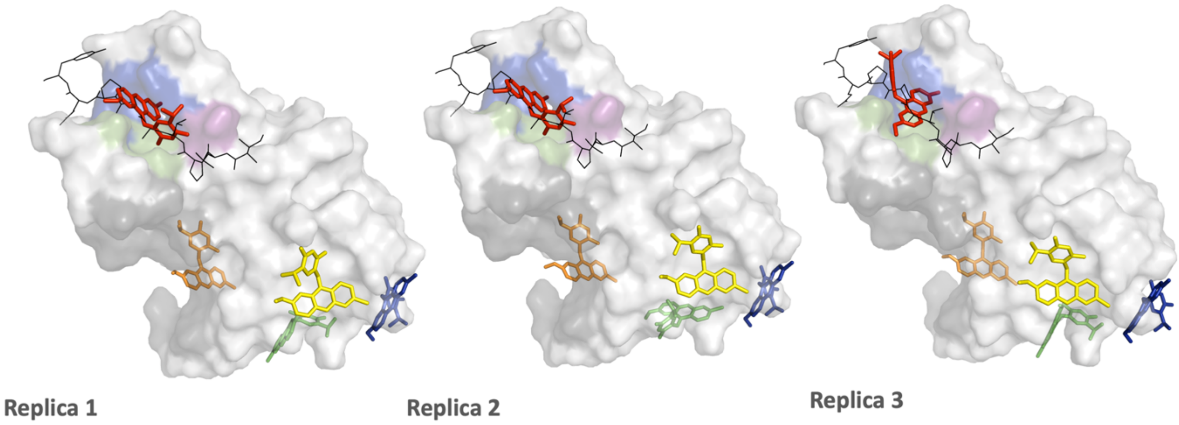


| UEV-10 | Replica 1 | Replica 2 | Replica 3 |
|---|---|---|---|
| Energy | Lead Finder / MM-PBSA (kJ·mol$^{-1}$) | | |
| Pose 1 | -36.6/-161 | -36.4/-193 | -37.7/-116 |
| Pose 2 | -33.6/-218 | -33.3/-210 | -33.6/-225 |
| Pose 3 | -33.0/-179 | -32.9/-157 | -32.3/-157 |
| Pose 4 | -30.3/-161 | -30.2/-170 | -29.3/-180 |
| Pose 5 | -29.3/-144 | -29.2/-130 | -29.2/-125 |

| | PTAP site | Replica 3-Pose 1 | Replica 1-Pose 1 | Replica 2-Pose 1 |
|---|---|---|---|---|
| **UEV-10** | **Energy LF/MMPBSA (kJ·mol$^{-1}$)** | **-37.7 / -116** | **-36.5/ -161** | **-36.3/ -193** |
| | Hydrophobic | Y63, R64, Y68, I70, P139, F142 | Y63, Y68, I70, P139, V141 | Y63, Y68, I70, P139, V141 |
| | Hbonds | Y68, S143 | N69, S143 | N69, S143 |
| | π-Stacking | Y68 | Y68 | Y68 |
|  | Halog-bond | ---- | F135 | F135 |
|  | **Ub site** | **----** | **----** | **---** |

**Figure S10.- Blind docking of UEV-10 compound onto TSG101-UEV.** Shown is the surface representation of the TSG101-UEV domain (white) in complex with the EBOV L-domain (4EJE.pdb). The PTAP and Ubiquitin binding sites are highlighted in color (blue: P0 pocket, green: A-1 pocket, magenta: T-2 pocket; grey: ubiquitin). The top five poses for UEV-10 from each independent replica are shown as colored sticks (Pose 1: red; Pose 2 orange; Pose 3: yellow; Pose 4: green; Pose 5: cyan). The peptide ligand corresponding to the EBOV L-domain in 4EJE is shown in black as a reference. The table summarizes the binding energies calculated by the Lead Finder and MM-PBSA algorithms together with the different forces and interacting residues contributing to binding for the best 3 poses in the PTAP and Ubiquitin sites.

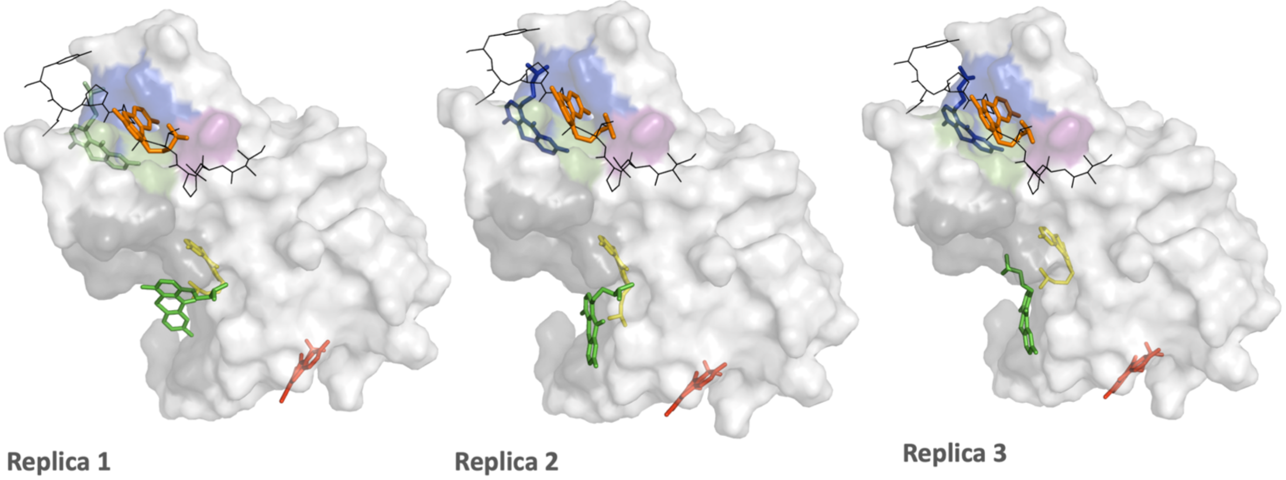


| UEV-11 | Replica 1 | Replica 2 | Replica 3 |
|---|---|---|---|
| Energy | Lead Finder / MM-PBSA (kJ·mol$^{-1}$) | | |
| Pose 1 | -35.9/-197 | -36.1/-191 | -36.5/-195 |
| Pose 2 | -34.9/-216 | -34.1/-223 | -34.9/-221 |
| Pose 3 | -32.1/-179 | -32.3/-182 | -32.4/-169 |
| Pose 4 | -29.5/-239 | -29.8/-224 | -29.0/-233 |
| Pose 5 | -28.0/-193 | -28.7/-167 | -29.0/-168 |

| | PTAP site | Replica 1-Pose 2 | Replica 3-Pose 2 | Replica 2-Pose 2 |
|---|---|---|---|---|
| | **Energy** LF/MMPBSA (kJ·mol$^{-1}$) | **-34.9 / -216** | **-34.9 / -221** | **-34.1 / -223** |
| | Hydrophobic | Y63, Y68, I70, P139 | Y63, Y68, I70, P139 | Y63, Y68, I70, P139 |
| | Hbonds | N69, F135 | N69, F135 | N69, F135 |
| **UEV-11** | π-Stacking | Y68 | Y68 | Y68 |
| | **Ub site** | **Replica 2-Pose 4** | **Replica 1-Pose 4** | **Replica 3-Pose 4** |
| | **Energy** LF/MMPBSA (kJ·mol$^{-1}$) | **-29.8 / -224** | **-29.4 / -239** | **-28.9 / -233** |
| | Hydrophobic | **Y42,** V43, **W75, F88** | **Y42,** K90 | **Y42,** V43, **F88** |
| | Hbonds | D40, K90 | V43, **W75,** K90 | D40, **W75** |
| | π-Stacking | | **F88** | **W75** |

**Figure S11.- Blind docking of UEV-11 compound onto TSG101-UEV.** Shown is the surface representation of the TSG101-UEV domain (white) in complex with the EBOV L-domain (4EJE.pdb). The PTAP and Ubiquitin binding sites are highlighted in color (blue: P0 pocket, green: A-1 pocket, magenta: T-2 pocket; grey: ubiquitin). The top five poses for UEV-11 from each independent replica are shown as colored sticks (Pose 1: red; Pose 2 orange; Pose 3: yellow; Pose 4: green; Pose 5: cyan). The peptide ligand corresponding to the EBOV L-domain in 4EJE is shown in black as a reference. The table summarizes the binding energies calculated by the Lead Finder and MM-PBSA algorithms together with the different forces and interacting residues contributing to binding for the best 3 poses in the PTAP and Ubiquitin sites.

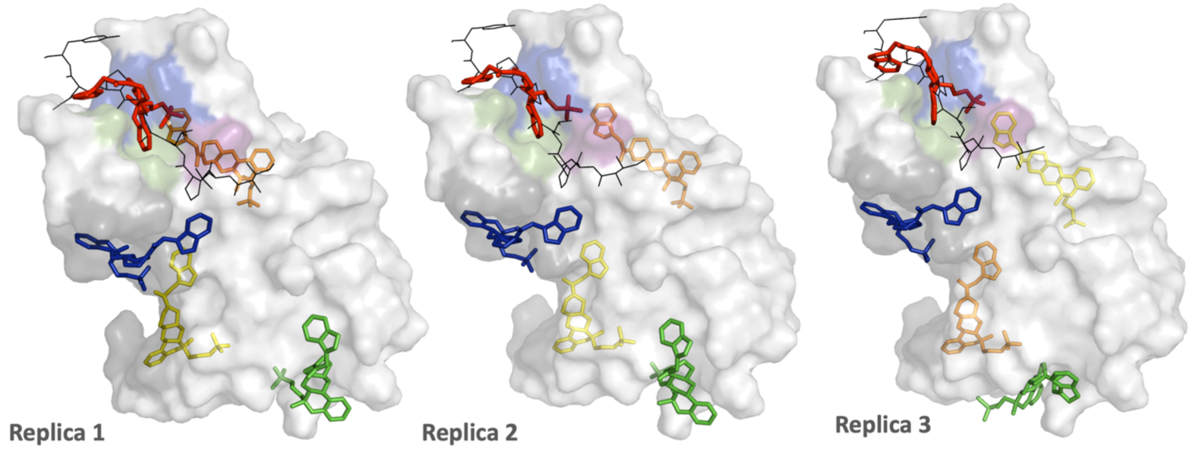


| UEV-21 | Replica 1 | Replica 2 | Replica 3 |
|---|---|---|---|
| **Energy** | **Lead Finder / MM-PBSA (kJ·mol$^{-1}$)** | | |
| Pose 1 | -37.9/-215 | -38.9/-201 | -38.7/-155 |
| Pose 2 | -37.8/-204 | -36.6/-230 | -36.9/-205 |
| Pose 3 | -36.6/-225 | -35.6/-250 | -36.7/-231 |
| Pose 4 | -34.2/-186 | -35.0/-203 | -36.4/-248 |
| Pose 5 | -33.2/-197 | -32.6/-232 | -32.8/-205 |

| | PTAP site | Replica 2-Pose1 | Replica 3-Pose1 | Replica 1-Pose1 |
|---|---|---|---|---|
| | Energy LF/MMPBSA (kJ·mol$^{-1}$) | **-38.9 / -201** | **-38.7 / -155** | **- 37.9/ -215** |
| | Hydrophobic | Y63, Y68, I70, P139, V141, F142 | V61, Y63, Y68, I70, P139, V141 | Y68, I70, P139, V141, F142 |
| | Hbonds | S143 | S143 | S143 |
| **UEV-21** | Halog-bond | V61 | | |
| | **Ub site** | **Replica 1-Pose5** | **Replica 3-Pose 5** | **Replica 2-Pose 5** |
| | Energy LF/MMPBSA (kJ·mol$^{-1}$) | **-33.2 / -197** | **-32.8 / -205** | **-32.6 / -232** |
| | Hydrophobic | T58, P71, **F88**, P91, T92, I97 | T58, P71, **F88**, T92, I97 | T58, P71, **F88**, K90, P91, T92, I97 |
| | Halog-bond | | V89 | |

**Figure S12.- Blind docking of UEV-21 compound onto TSG101-UEV.** Shown is the surface representation of the TSG101-UEV domain (white) in complex with the EBOV L-domain (4EJE.pdb). The PTAP and Ubiquitin binding sites are highlighted in color (blue: P0 pocket, green: A-1 pocket, magenta: T-2 pocket; grey: ubiquitin). The top five poses for UEV-21 from each independent replica are shown as colored sticks (Pose 1: red; Pose 2 orange; Pose 3: yellow; Pose 4: green; Pose 5: cyan). The peptide ligand corresponding to the EBOV L-domain in 4EJE is shown in black as a reference. The table summarizes the binding energies calculated by the Lead Finder and MM-PBSA algorithms together with the different forces and interacting residues contributing to binding for the best 3 poses in the PTAP and Ubiquitin sites.

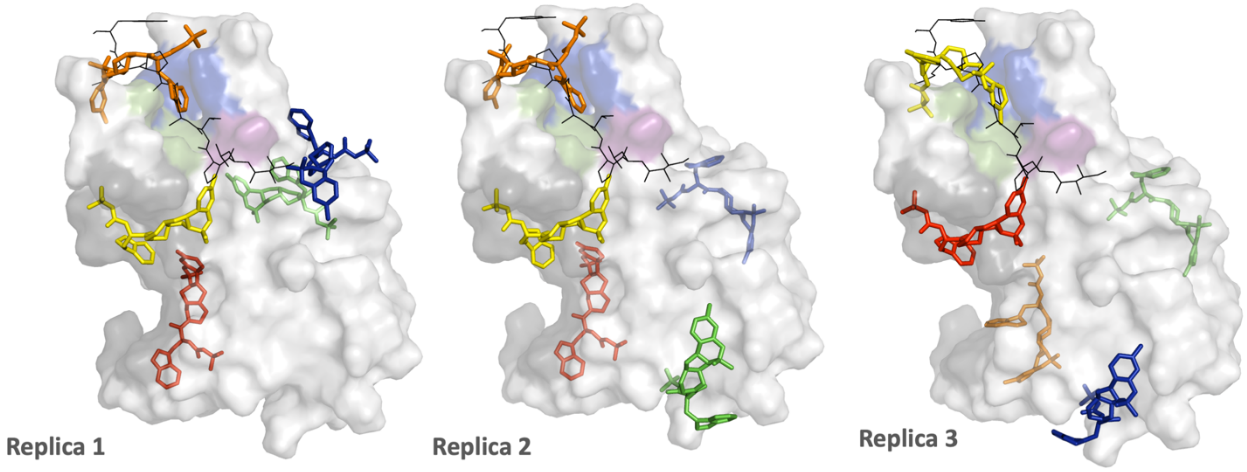


| UEV-22 | Replica 1 | Replica 2 | Replica 3 |
|---|---|---|---|
| **Energy** | **Lead Finder / MM-PBSA (kJ·mol$^{-1}$)** | | |
| Pose 1 | -37.8/-202 | -36.9/-208 | -36.7/200 |
| Pose 2 | -34.4/-216 | -36.9/-158 | -35.4/-222 |
| Pose 3 | -34.1/-199 | -36.2/-208 | -35.1/-178 |
| Pose 4 | -32.7/-200 | -35.9/-161 | -34.1/-177 |
| Pose 5 | -30.3/-167 | -33.1/-165 | -31.2/-192 |

| | **PTAP site** | **Replica 1-Pose 2** | **Replica 2-Pose 2** | **Replica 3-Pose 3** |
|---|---|---|---|---|
| **UEV-22** | **Energy LF/MMPBSA (kJ·mol$^{-1}$)** | **-34.4 / -216** | **-36.8 / -158** | **-35.1 / -178** |
| | Hydrophobic | Y63, R64, I70, K98, P139, F142 | Y63, Y68, I70, K98, P139, F142 | Y63, Y68, I70, V141, F142, P145 |
| | Hbonds | Y68, S143 | Y68, S143 | S143 |
| | π-Stacking | Y68 | Y68 | Y68 |
| | **Ub site** | **Replica 3-Pose 1** | **Replica 2-Pose 3** | **Replica 1-Pose 3** |
| | **Energy LF/MMPBSA (kJ·mol$^{-1}$)** | **-36.7 / -200** | **-36.2 /-208** | **-34.1 / -199** |
| | Hydrophobic | T58, **F88**, K90, T92 | T58, P71, **F88**, K90, T92 | T58, P71, **F88**, K90, T92 |
| | Hbonds | P91 | V89, P91 | P91, N106 |
| | Cation-π | | K90 | |

**Figure S13.- Blind docking of UEV-22 compound onto TSG101-UEV.** Shown is the surface representation of the TSG101-UEV domain (white) in complex with the EBOV L-domain (4EJE.pdb). The PTAP and Ubiquitin binding sites are highlighted in color (blue: P0 pocket, green: A-1 pocket, magenta: T-2 pocket; grey: ubiquitin). The top five poses for UEV-22 from each independent replica are shown as colored sticks (Pose 1: red; Pose 2 orange; Pose 3: yellow; Pose 4: green; Pose 5: cyan). The peptide ligand corresponding to the EBOV L-domain in 4EJE is shown in black as a reference. The table summarizes the binding energies calculated by the Lead Finder and MM-PBSA algorithms together with the different forces and interacting residues contributing to binding for the best 3 poses in the PTAP and Ubiquitin sites.

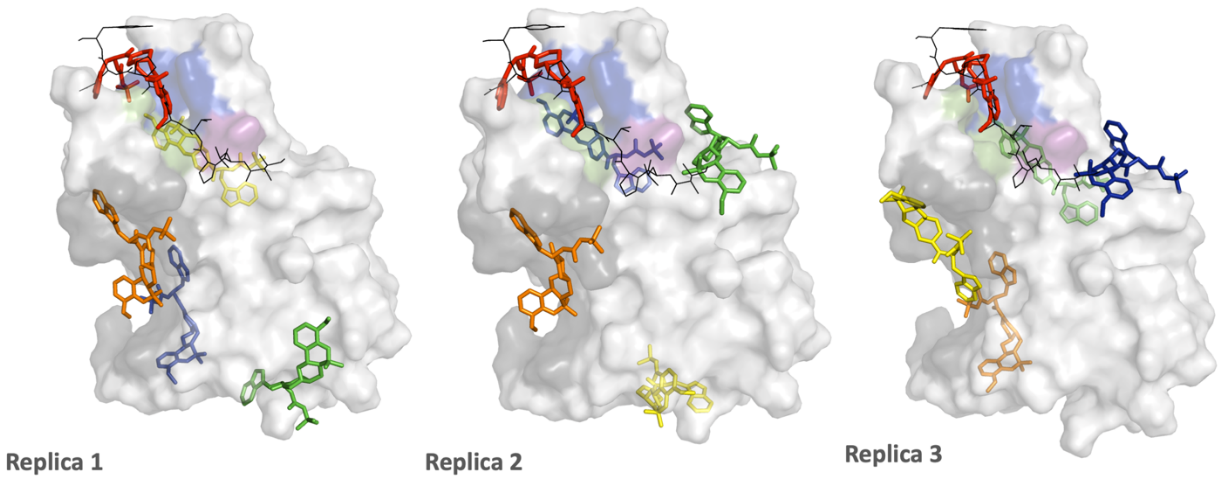


| UEV-23 | Replica 1 | Replica 2 | Replica 3 |
|---|---|---|---|
| Energy | Lead Finder / MM-PBSA (kJ·mol$^{-1}$) | | |
| Pose 1 | -37.8/-267 | -37.3/-257 | -37.6/-234 |
| Pose 2 | -34.4/-280 | -36.1/-190 | -36.1/-290 |
| Pose 3 | -34.1/-222 | -34.4/-191 | -34.4/-176 |
| Pose 4 | -32.7/-208 | -32.8/-189 | -32.8/-253 |
| Pose 5 | -30.3/-195 | -32.5/-234 | -32.0/-182 |

| | PTAP site | Replica 1-Pose 1 | Replica 3-Pose1 | Replica 2-Pose1 |
|---|---|---|---|---|
| UEV-23 | Energy LF/MMPBSA (kJ·mol$^{-1}$) | **-37.7 / -267** | **-37.6 / -234** | **-37.2 / -257** |
| | Hydrophobic | Y63, Y68, I70, P139, F142 | Y63; Y68, F142 | Y63, Y68, E138, F142 |
| | Hbonds | Y63, K98, S143 | Y63, K98, S143 | Y63, K98, S143 |
| | π-Stacking | Y68, F142 | Y68, F142 | Y68, F142 |
| | **Ub site** | **Replica 2-Pose 2** | **Replica 1-Pose 2** | **Replica 3-Pose 3** |
| | Energy LF/MMPBSA (kJ·mol$^{-1}$) | **-36.1 / -190** | **-36.0 / -290** | **-34.4 / -176** |
| | Hydrophobic | **Y42, W75, F88,** P91 | **Y42, W75, F88**, P91, T92 | **Y42, W75, K90**, I97 |
| | Hbonds | K90 | P91, T92 | **W75,** R144 |
| | π-Stacking | | | **F88** |

**Figure S14.- Blind docking of UEV-23 compound onto TSG101-UEV.** Shown is the surface representation of the TSG101-UEV domain (white) in complex with the EBOV L-domain (4EJE.pdb). The PTAP and Ubiquitin binding sites are highlighted in color (blue: P0 pocket, green: A-1 pocket, magenta: T-2 pocket; grey: ubiquitin). The top five poses for UEV-23 from each independent replica are shown as colored sticks (Pose 1: red; Pose 2 orange; Pose 3: yellow; Pose 4: green; Pose 5: cyan). The peptide ligand corresponding to the EBOV L-domain in 4EJE is shown in black as a reference. The table summarizes the binding energies calculated by the Lead Finder and MM-PBSA algorithms together with the different forces and interacting residues contributing to binding for the best 3 poses in the PTAP and Ubiquitin sites. .

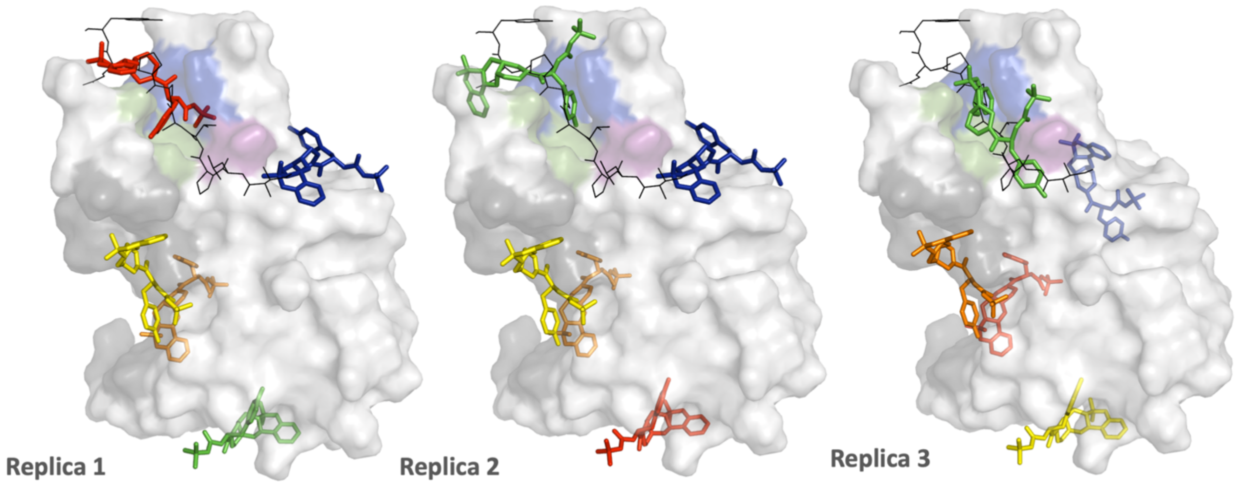


| UEV-24 | Replica 1 | Replica 2 | Replica 3 |
|---|---|---|---|
| Energy | Lead Finder / MM-PBSA (kJ·mol$^{-1}$) | | |
| Pose 1 | -38.8/-246 | -37/-191 | -37.8/-252 |
| Pose 2 | -38.5/-275 | -36.8/-265 | -36.9/-284 |
| Pose 3 | -36.6/-231 | -36.7/-231 | -36.6/-138 |
| Pose 4 | -36.3/-178 | -35.8/-179 | -35/-283 |
| Pose 5 | -32.6/-190 | -33.1/-196 | -31.4/-175 |

| | PTAP site | Replica 1-Pose 1 | Replica 2-Pose 4 | Replica 3-Pose 4 |
|---|---|---|---|---|
| UEV-24 | **Energy** LF/MMPBSA (kJ·mol$^{-1}$) | **-38.8 / -246** | **-35.8 / -179** | **-35.0 / -283** |
| | Hydrophobic | Y63, Y68, I70, M95, P139, F142, P145 | Y63, Y68, I70, P139, F142, P145 | T58, Y63, Y68, N69, I70, P139 |
| | Hbonds | S94 | V141, S143 | D34, N69, S143 |
| | π-Stacking | | Y68, F142 | Y68 |
| | **Ub site** | **Replica 3-Pose 2** | **Replica 2-Pose 3** | **Replica 1-Pose 3** |
| | **Energy** LF/MMPBSA (kJ·mol$^{-1}$) | **-36.8 / -284** | **-36.7 / -231** | **-36.6 / -231** |
| | Hydrophobic | **Y42, F88**, K90, T92 | D40, **Y42**, K90, T92 | **Y42, F88**, K90, T92 |
| | Hbonds | S41, V43, **W75**, K90 | S41, V43, **W75**, K90 | S41, V43, **W75**, K90 |

**Figure S15.- Blind docking of UEV-24 compound onto TSG101-UEV.** Shown is the surface representation of the TSG101-UEV domain (white) in complex with the EBOV L-domain (4EJE.pdb). The PTAP and Ubiquitin binding sites are highlighted in color (blue: P0 pocket, green: A-1 pocket, magenta: T-2 pocket; grey: ubiquitin). The top five poses for UEV-24 from each independent replica are shown as colored sticks (Pose 1: red; Pose 2 orange; Pose 3: yellow; Pose 4: green; Pose 5: cyan). The peptide ligand corresponding to the EBOV L-domain in 4EJE is shown in black as a reference. The table summarizes the binding energies calculated by the Lead Finder and MM-PBSA algorithms together with the different forces and interacting residues contributing to binding for the best 3 poses in the PTAP and Ubiquitin sites.
